# Phytochromes measure photoperiod in Brachypodium

**DOI:** 10.1101/697169

**Authors:** Mingjun Gao, Feng Geng, Cornelia Klose, Anne-Marie Staudt, He Huang, Duy Nguyen, Hui Lan, Todd C. Mockler, Dmitri A. Nusinow, Andreas Hiltbrunner, Eberhard Schäfer, Philip A. Wigge, Katja E. Jaeger

**Affiliations:** Sainsbury Laboratory, University of Cambridge, 47 Bateman St., Cambridge CB2 1LR, UK.; Institut für Biologie II, University of Freiburg, Schaenzlestr. 1, D-79104 Freiburg, Germany.; Donald Danforth Plant Science Center, St. Louis, MO 63132, USA; Signalling Research Centres BIOSS and CIBSS, University of Freiburg, Schaenzlestr. 18, 79104 Freiburg, Germany; BIOSS Centre for Biological Signalling Studies, University of Freiburg, Schaenzlestr. 18, 79104 Freiburg, Germany; Leibniz-Institut für Gemüse- und Zierpflanzenbau, Theodor-Echtermeyer-Weg 1, 14979 Großbeeren, Germany; Institute of Biochemistry and Biology, University of Potsdam, 14476 Potsdam, Germany

## Abstract

Daylength is a key seasonal cue for animals and plants. In cereals, photoperiodic responses are a major adaptive trait, and alleles of clock genes such as *PHOTOPERIOD DEPENDENT1 (PPD1)* and *EARLY FLOWERING3 (ELF3)* have been selected for in breeding barley and wheat for more northern latitudes (Faure et al., 2012; Turner, Beales, Faure, Dunford, & Laurie, 2005). How monocot plants sense photoperiod and integrate this information into growth and development is not well understood. We show that in *Brachypodium distachyon*, phytochrome C (phyC) acts as a molecular timer, directly communicating information to the circadian clock protein ELF3. In this way, ELF3 levels integrate night length information. ELF3 is a central regulator of photoperiodism in Brachypodium, and *elf3* mutants display a constitutive long day transcriptome. Conversely, conditions that result in higher levels of ELF3 suppress long day responses. We are able to show that these effects are direct, as ELF3 and phyC occur in a common complex, and they associate with the promoters of a number of conserved regulators of photoperiodism, including *PPD1*. Consistent with observations in barley, we are able to show that *PPD1* overexpression accelerates flowering in SD and is necessary for rapid flowering in response to LD. These findings provide a conceptual framework for understanding observations in the photoperiodic responses of key crops, including wheat, barley and rice.

## Introduction

Flowering is a major developmental transition, and plants have evolved pathways to flower in response to seasonal cues to maximise their reproductive fitness (Song, Shim, Kinmonth-Schultz, & Imaizumi, 2015). Photoperiod provides a key seasonal cue, and in temperate climates, long photoperiods serve as a signal of spring and summer, accelerating flowering in many plants. In *Arabidopsis thaliana*, long days result in the stabilization of the floral activator CONSTANS (CO), which activates the expression of the florigen encoding gene *FLOWERING LOCUS T (FT)* (Hayama et al., 2017). Temperate grasses, such as Brachypodium, barley and wheat also induce flowering through the induction of *FT*-related genes, however there are differences in the signalling pathways activating *FT* expression.

The major regulator of natural variation in photoperiod responsiveness in barley is the transcriptional regulator *PHOTOPERIOD DEPENDENT1* (Hv-Ppd1), first identified as dominant allele that accelerates flowering under short day conditions, making plants photoperiod insensitive (Turner et al., 2005). Analyses of *ppd1* alleles indicate that promoter insertions and deletions have played a major role modulating *PPD1* expression, revealing a 95 bp region within the promoter that is conserved between wheat, barley and Brachypodium (Seki et al., 2013; Wilhelm, Turner, & Laurie, 2009). It has been hypothesized that a photoperiod dependent repressor may bind this 95 bp region short days to inhibit flowering. *Ppd-H1* also influences leaf size, a trait which is under photoperiod control, consistent with *Ppd-H1* being a key output of the photoperiod pathway in grasses (Digel et al., 2016). The evening complex (EC), an integral component of the circadian clock, is also a key regulator of photoperiodism in grasses. The *early maturity8 (eam8)* allele in barley confers early flowering in SD, and encodes the barley ortholog of *EARLY FLOWERING3 (ELF3)* (Faure et al., 2012), and in wheat, *Earliness Per Se (eps)* also confers early flowering and is likely caused by mutation in an *ELF3* related gene (Alvarez, Tranquilli, Lewis, Kippes, & Dubcovsky, 2016). Similarly, *eam10* encodes HvLUX, and is necessary for correctly responding to photoperiod (Campoli et al., 2013), while *PHYTOCLOCK (LUX)* alleles also confer early flowering in wheat (Mizuno et al., 2016).

Unlike in Arabidopsis, where phytochromes repress flowering, PhyC is an essential inducer of flowering in Brachypodium (Woods, Ream, Minevich, Hobert, & Amasino, 2014), and interfering with phyC in barley and wheat also greatly delays flowering, indicating that phyC is an essential input for photoperiodism (Chen et al., 2014; Nishida et al., 2013). In barley, the allele *eam5*, which shows constitutively early flowering, is associated with a mutation in the GAF domain of phyC. Since the GAF domains of phytochromes influence their behaviour, this suggests that the interconversion of phyC between the Pfr and Pr states may be important in floral regulation (Pankin et al., 2014). Finally, *phyC-1* in Brachypodium also shows additional photoperiod phenotypes such as leaf morphology differences as well as flowering time (Woods et al., 2014) indicating a major role for phytochrome signalling, not just in flowering but photoperiod responses in general. How the EC and *PPD1* influence flowering, and how phyC conveys photoperiod information to these regulators is however not well understood.

## Results

To determine if the role of *ELF3* in flowering is conserved in Brachypodium, we created loss of function alleles in *ELF3*. *elf3-1* plants show constitutive early flowering, largely independent of photoperiod, indicating that *ELF3* is necessary for responding to photoperiod (**Fig. 1A-C**). We find that *ELF3* overexpressing plants show delayed flowering in long days, suggesting that *ELF3* is necessary and sufficient to transmit photoperiodic signals in Brachypodium (**Fig. 1D** and **E**). To understand how *ELF3* may be controlling photoperiodic responses in Brachypodium we performed affinity purification coupled with mass spectrometry to identify the ELF3 protein interactome. Consistent with the evening complex being conserved between Arabidopsis and monocots (Huang et al., 2017), ELF4 and LUX are detected as ELF3 interactors (**Fig. 1F; Supplementary Dataset S1**). The identification of two TOPLESS (TPL)-related proteins suggests a mechanism by which the evening complex represses gene expression. Photoperiodism in Arabidopsis is also mediated by the repression of *FT* and *CO* by a TPL containing transcriptional complex, indicating this may be a common mechanism to achieve photoperiodic gene expression (Goralogia et al., 2017). Photoperiodism requires light perception, and we identified the light sensing phytochromes phyB and phyC as ELF3 interactors. Brachypodium contains three phytochromes, and we therefore investigated the extent to which phytochromes are necessary for photoperiodism. *phyC-4* does not flower under LD, consistent with previous reports (**Fig. S1A** and **B**) (Woods et al., 2014), while *phyA-1* and *phyB-1* both show delayed responses to LD (**Fig. S1C-F**). These results suggest that phytochromes act in the same pathway as *ELF3*.

**Fig. 1.**
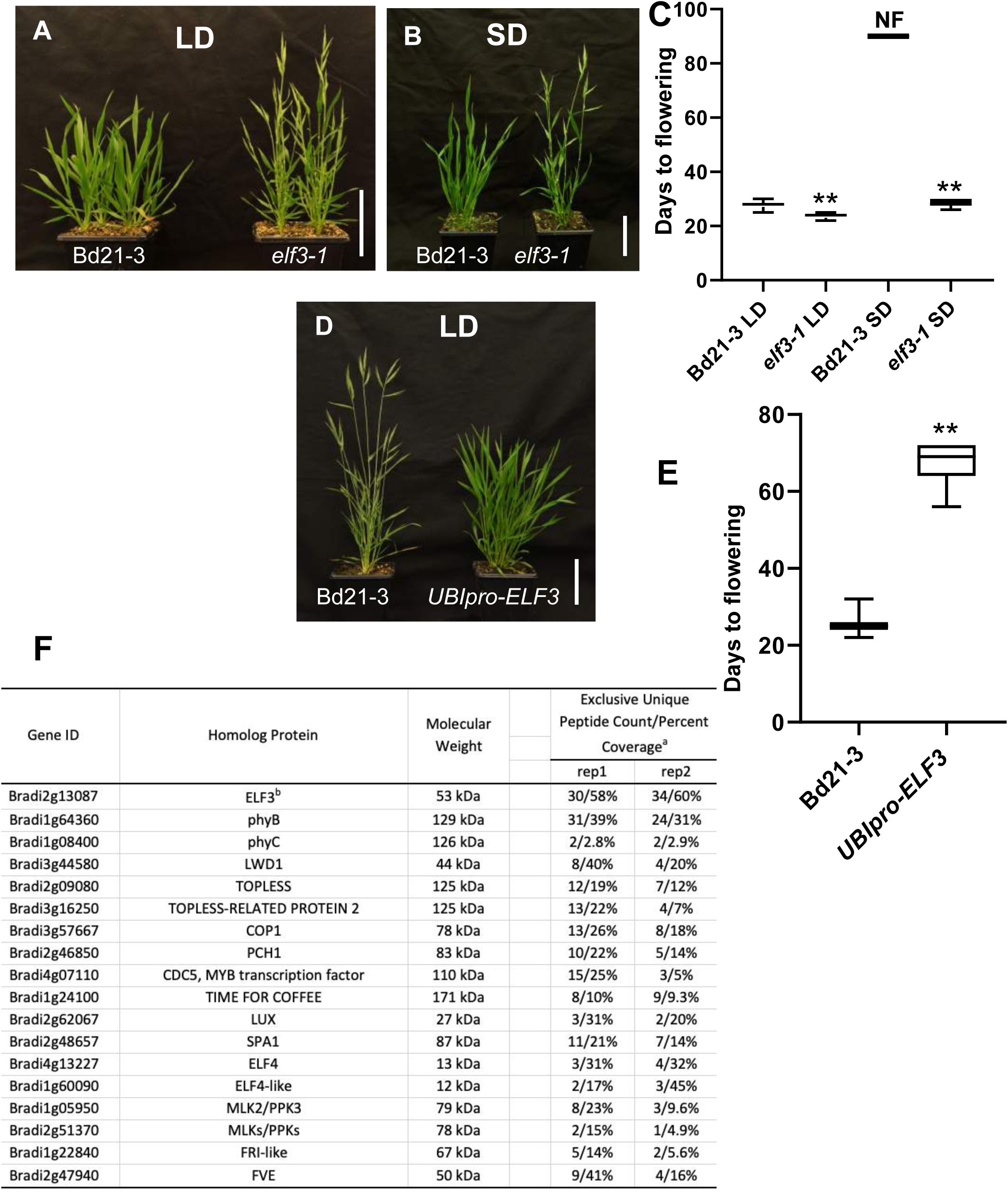
*ELF3* is necessary for photoperiodism in *Brachypodium.* (**A** to **C**) *elf3-1* shows a constitutive long day flowering phenotype under short day conditions, where wild-type does not flower (NF). (Student’s t-test, **P < 0.01). (**D** to **E**) Constitutive expression of *ELF3* under the *UBIQUITIN* promoter (*UBI_pro_)* is sufficient to greatly delay flowering under inductive long day conditions. (Student’s t-test, **P < 0.01). (**F**) Curated list of proteins binding ELF3 identified by mass spectrometry. No peptides for these proteins detected in the negative control (YFP-HFC). ^a^All proteins match 99% threshold with minimum 2 peptides

Flowering is a complex trait, and it is possible that the perturbation of the floral transition we observe in *phyC-4* and *elf3-1* is indirect. We therefore compared the transcriptomes of *elf3-1* and *phyC-4* with that of wild-type plants in response to LD (**Fig. 2**, **Fig. S2**, **Fig. S3** and **Supplementary dataset S2**). A large number of genes are induced in the afternoon and evening by LD in two-week old seedlings, and multiple clusters show enhanced expression in response to photoperiod (particularly 8, 13, 19, 22 and 26). These LD expressed clusters are also induced in two-week old *elf3-1* seedlings grown in SD, indicating that *ELF3* is necessary for photoperiod dependent expression of the transcriptome. This is also apparent when comparing the *elf3-1* transcriptome under LD with *elf3-1* under SD, here there is very little difference in expression, indicating that *elf3-1* has a photoperiod insensitive transcriptome. The constitutive early flowering behaviour of *elf3-1* plants is therefore reflected in these plants having a constitutive LD transcriptome even when grown under SD. (**Fig. S3**). Two clusters (4 and 34) appear to reflect mainly developmental age, and they are primarily differentially expressed in *elf3-1* at week 3 compared to week 2. Cluster 34 contains *FT1, FT2, BdAP1* and *BdMAD5*, all indicators of flowering, consistent with these clusters containing downstream targets (**Supplementary dataset S3**). In contrast to *elf3-1*, *phyC-4* plants show the opposite phenotype, with large scale repression of those genes that are induced by LD in wild-type (**Fig. 2** and **Fig. S3**). Many of the photoperiod response clusters, which are differentially affected by *elf3-1* and *phyC-4* contain classical photoperiod and circadian genes. Cluster 8 displays a characteristic evening complex target gene behaviour, being more expressed in *elf3-1* at the end of the day. This cluster contains classical clock genes such as *GI, PPD1* and *BdLNK1*, and is enriched for genes involved in photosynthesis, as has been observed for EC targets in Arabidopsis (**Fig. S4**) (Ezer et al., 2017). Cluster 22 shows a strong photoperiod response, and *elf3-1* mimics this pattern of expression with repression in the morning, while *phyC-4* shows the opposite behaviour. This cluster has photoperiod and clock-related genes including *PIF3, CCA1, RVE6* and *RVE7*. These results indicate that *ELF3* is necessary for repressing the LD transcriptome while *PHYC* is required for activating gene expression in response to inductive photoperiods.

**Fig. 2.**
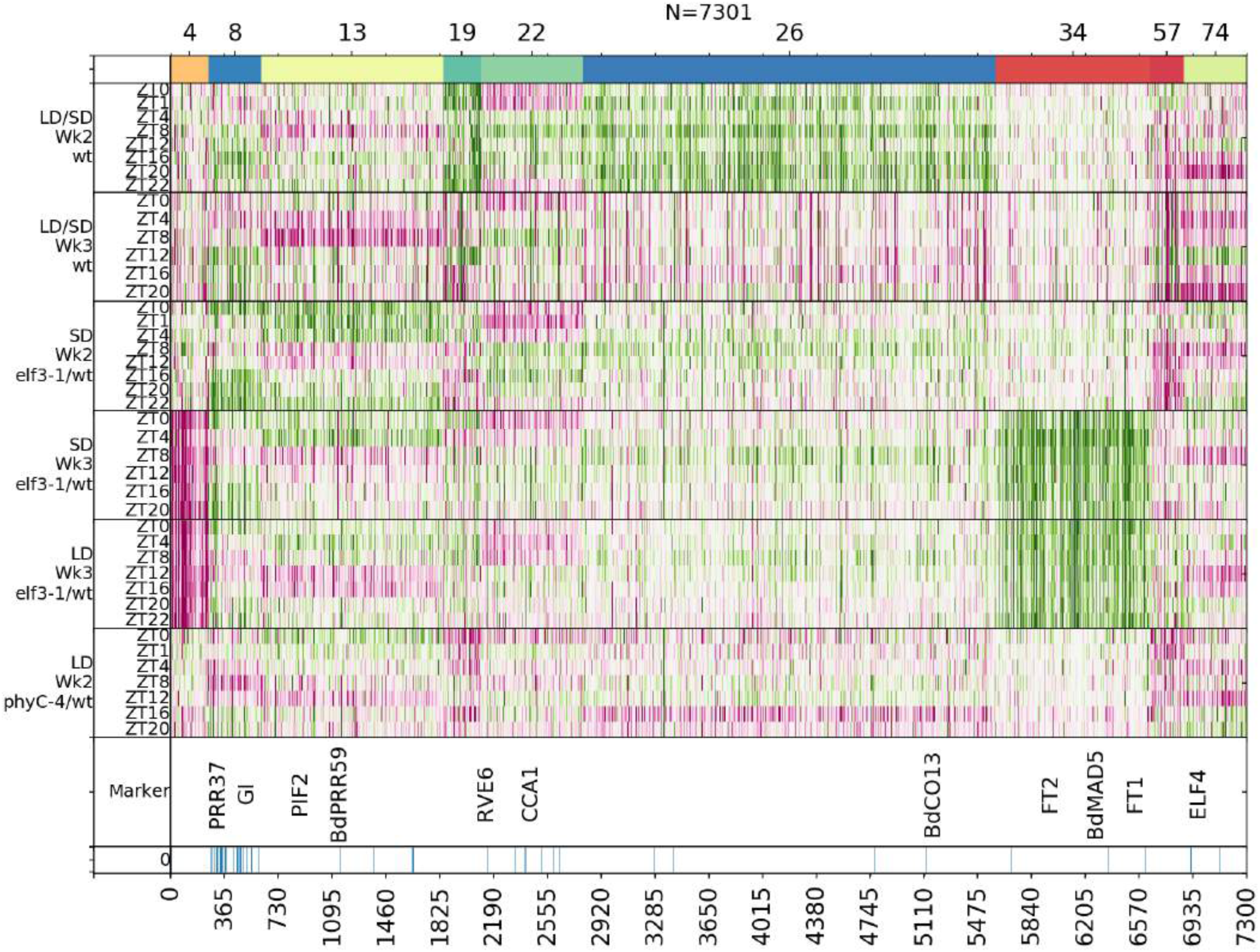
*elf3-1* displays a constitutive LD transcriptome and *phyC-4* resembles a SD grown plant. Clustering by gene expression reveals that many of the clusters of genes that respond to LD show a similar behaviour in *elf3-1* grown in SD, suggesting that this background has a constitutive LD response. *phyC-4* shows the opposite response, resembling a SD plant when grown under LD. Clusters 8, 13, 19, 22 and 26 in particular show up-regulation in response to LD, they are expressed in response to the *elf3-1* mutation, and down-regulated in *phyC-4*. Clusters 4 and 34 appear to reflect developmental changes in response to the floral transition, reflecting the very early flowering phenotype of *elf3-1*.

To understand how ELF3 influences photoperiodic gene expression we identified ELF3 target genes by ChIP-seq. Consistent with what has been seen in Arabidopsis, we observe binding of ELF3 at the promoters of clock genes such as *GIGANTEA (GI)* (**Fig. 3A**). We observe that the EC appears to be photoperiod responsive since ELF3 occupancy on promoters is strongly affected by the time of day and the light regime. Binding is maximal in short days during and at the end of the night, but rapidly declines in response to light. Continuous light or LD greatly reduce ELF3 occupancy (**Fig. 3B**). We used this dynamic change in ELF3 occupancy as a signature to identify putative biologically relevant ELF3 ChIP-seq peaks. In this way, we identified 94 genes associated with time of day responsive ELF3 ChIP-seq peaks (**Supplementary Dataset S4**). About 1/3 of these genes associated with ELF3 ChIP-seq peaks show an *ELF3* dependent expression pattern (**Fig. 3C**). In Arabidopsis ELF3 functions as a transcriptonal repressor (Ezer et al., 2017), and we observe the same behaviour in Brachypodium, as ELF3 bound genes are strongly up-regulated at the end of the day in *elf3-1* (**Fig. S5**). Many genes bound by ELF3 are induced by LD as well as *elf3-1*, while they are repressed in *phyC-4* (**Fig. 3D** and **Fig. S6**), consistent with ELF3 being a central regulator of photoperiod controlled gene expression.

**Fig. 3.**
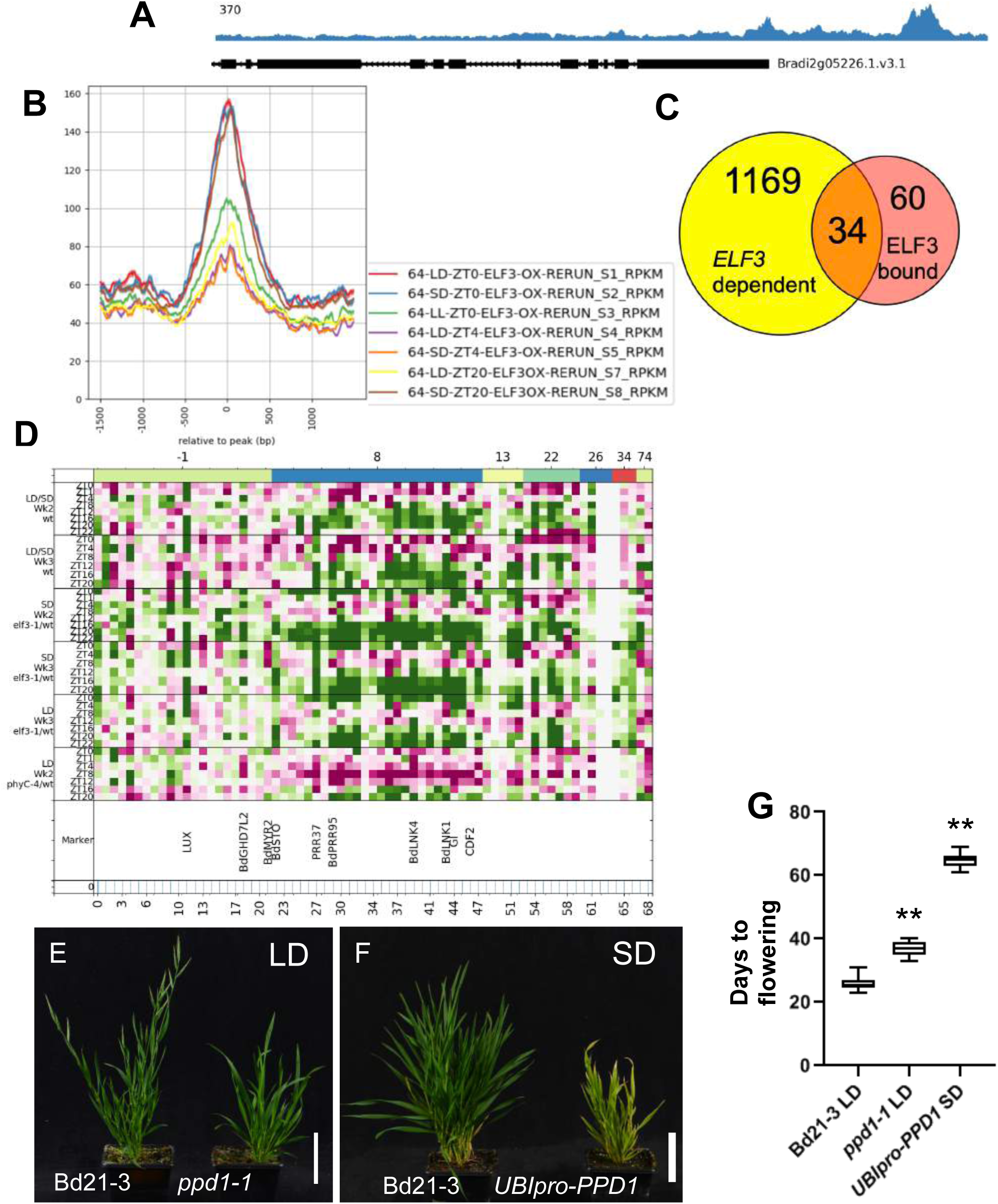
ELF3 directly controls many photoperiod responsive genes. **(A)** ELF3 associates with the promoter of *GI* as measured by ChIP-seq. (B) The occupancy of ELF3 on target promoters responds to photoperiod. Binding increases in short days at ZT0 and ZT20, but sharply decreases at ZT4. This is a response to light, since binding at ZT0 and ZT20 is sharply reduced in LD and LL. (C) There are 94 genes differentially bound by ELF3 and about 1/3 of these genes have perturbed expression in *elf3-1 (D) T*hese differentially bound direct targets of ELF3 show a high degree of responsiveness to photoperiod. Cluster −1 collects genes without a confident cluster. Genes which are upregulated are shown in green, genes which are downregulated are shown in red and unchanged is depicted in white. (E-G) *ppd1-1* is required for photoperiodic acceleration of flowering and UBIpro-PPD1 flower independently of photoperiod

Flowering is controlled by *FT* (florigen) genes (Wigge, 2011), and we find that *FT1* shows the strongest response and absolute expression level in response to inductive photoperiods. Ectopic overexpression of *FT1* is sufficient to restore flowering in SD (**Fig. S7**) suggesting the induction of this gene in response to long photoperiod is important in the activation of flowering. Although we see some evidence for binding of ELF3 at the promoter of *FT2* (**Fig. S8**), no binding at *FT1* is observed, suggesting that ELF3 transmits photoperiodism signals to *FT1* indirectly.

To identify potential ELF3 bound genes that may act as activators of *FT1*, we searched for transcripts encoding transcriptonal regulators induced by LD, age and *elf3-1* and repressed in *phyC-4*. *PPD1/PRR37* shows a particularly strong response, and we observe a correlation of *FT1* and *PPD1* expression with age (**Fig. S6** and **S9**). The binding of ELF3 to the *PPD1* promoter is also responsive to photoperiod (**Fig. S10** and **11**). Consistent with this apparent role for ELF3 in controlling *PPD1/PRR37* expression, *ELF3-OX* plants show a strong down-regulation of *PPD1/PRR37* transcript levels, which is accompanied by a loss of *FT1* expression (**Fig. S12**).

By genome-editing we created a loss of function allele, *ppd1-1*, which we confirmed as being late flowering in LD, while expression of *PPD1* under the *UBIQUITIN* promoter (*PPD1-OX*) is sufficient to activate flowering in SD (**Fig. 3E-G**). Consistent with this phenotype, we observe a strong up-regulation of *FT1* in *PPD1-OX*, as well as that of *FT2* and *AP1* (**Fig. S13**). Altering *PPD1* activity also affects a number of photoperiod responsive transcripts, but not as many as seen for *elf3-1* and *phyC-4*, consistent with *PPD1/PRR37* being downstream of these genes in the photoperiod pathway (**Fig. S6**). Since *ppd1-1* does not have as strong a flowering phenotype as *phyC-4* this indicates that other genes contribute to the activation of *FT1* in parallel to *PPD1*. Possible candidates include four genes encoding B-box Proteins (BBX), including the photoperiod regulator *CONSTANS (CO)* as well as the Brachypodium gene related to *GHD7* that controls flowering in rice. Additionally, 4 Cycling DOF (*CDF*) encoding genes are also directly bound by ELF3 (**Supplementary Table S4**).

Taken together, these results suggest a model whereby ELF3 represses the expression of *PPD1* and other flowering regulators, and under inductive photoperiods ELF3 levels decline, enabling the up-regulation of these transcriptional activators and the activation of *FT* genes and flowering. The expression of *ELF3* however is largely constant and does not show significant circadian variation, and it is expressed in both SD and LD and is unchanged in *phyC-4* compared to wild-type (**Fig. S14**). This suggests that the regulation of *ELF3* may be post-translational. Consistent with this hypothesis, the late flowering phenotype of *ELF3-OX* is sensitive to light exposure (**Fig. 4A** and **B**). While *ELF3-OX* plants are very late flowering in LD, continuous light (LL) is able to accelerate flowering, consistent with our observations for the degree of binding of ELF3 to target promoters (**Fig. 3B**; **Fig. S10** and **S11**). We therefore hypothesized that ELF3 protein levels are light sensitive. Indeed, ELF3 protein accumulates at the end of the night to high levels under SD, and is rapidly degraded upon exposure to light. A similar pattern is seen under LD, but the levels of ELF3 are lower (**Fig. 4C** and **D**, **Fig. S15**).

**Fig. 4.**
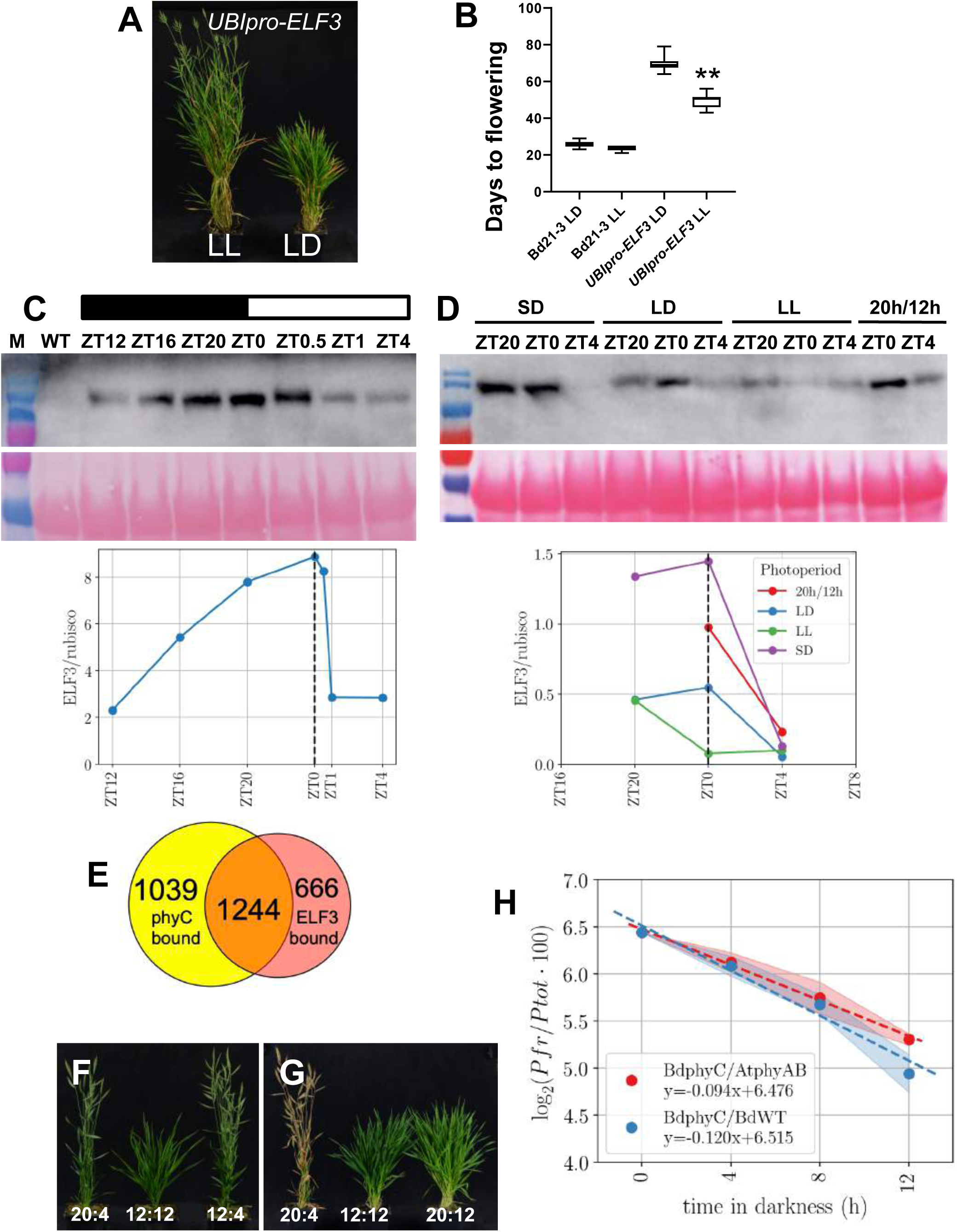
ELF3 protein levels integrate photoperiod information. (**A** to **B**) The late flowering phenotype of *UBIpro-ELF3* in long days is partially suppressed by growth in continuous light. (**D** and **D**) ELF3 protein levels accumulate during the night and are rapidly reduced on exposure to light. (**E**) Overlap between phyC and ELF3 ChIP-seq peaks. (**F**) phyC dark reversion has a half-life of about 8 h. **G** and **H**) Night length but not day length is the key determinant of when plants will flower.

Since *phyC-4* transcriptionally resembles a plant with elevated ELF3 signaling, this suggests that phyC may be the major light receptor controlling ELF3 activity. To determine if this occurs via a direct mechanism, we performed ChIP-seq of phyC. In Arabidopsis, phyB binds to target genes to modulate their expression (Jung et al., 2016), and we investigated if this might be true for phyC in Brachypodium. We observe coincidence between ELF3 and phyC ChIP-seq peaks for many key genes such as *LUX* (**Fig. 4E** and **Fig. S16**; **Supplementary Dataset S5**). Phytochromes have been observed to interact with ELF3 in other systems, so this finding is consistent with these observations and suggests phyC may counteract the ability of ELF3 to repress its targets.

In Arabidopsis, photoperiod is measured through the interaction of phototransduction pathways with the circadian rhythm, and flowering is activated when light corresponds with a sensitive phase of the circadian cycle (Song et al., 2015). This has been demonstrated using non-24 h light-dark cycles (T-cycles), an approach first described by Nanda and Hamner (Nanda & Hamner, 1958). Flowering does not correlate with the length of the light or dark for non-24 h T-cycles, indicating that Arabidopsis does not measure night or day length alone to determine photoperiodism (Roden, Song, Jackson, Morris, & Carre, 2002). By contrast, other plants such as *Xanthium* measure night length to determine photoperiodism (Hamner, 1940; Thomas & Vince-Prue, 1997). Our results so far indicate that night-length measurement via the integration of Pfr levels is an important mechanism in Brachypodium. This predicts that, unlike for Arabidopsis, modulating night-length alone will have the largest effect on flowering in Brachypodium. To test this, we grew plants under a range of T-cycles. As expected SD control plants under 12L:12D do not flower, while LD (20L:4D) plants do (**Fig. 4F** and **G** and **Fig. S17**). Combining a long day with a long night (20L:12D) also prevents flowering, consistent with night-length being the main determinant of the photoperiodic response in Brachypodium. A long day is not required for flowering, since plants grown under 12L:4D cycles are early flowering. In Arabidopsis, growing plants under different T-cycles alters the phase of the circadian rhythm with the light-dark cycle. For example plants grown in 10L:20D actually flower earlier than plants grown under 10L:14D, although they both experience the same duration of day length. To investigate whether some of the effects we observe might be due to alterations in the phasing of the external day-night cycle with endogenous rhythms, we grew plants under 12L:20D, extending the usual night by 8 h and thereby altering the phasing of the circadian clock compared to 12L:12D grown plants. These plants also display a SD phenotype and do not flower, suggesting that absolute night-length plays a major role in determining photoperiodic responses in Brachypodium (**Fig. S17**).

Since phytochromes respond to light rapidly, we predicted that brief exposure to light during the dark period (“night-break”) would be sufficient to overcome the repressive effects of long nights on flowering. In the case of rice, the introduction of a night-break prevents flowering (Ishikawa, Shinomura, Takano, & Shimamoto, 2009), while in wheat flowering can be activated by night-break (Pearce et al., 2017). Consistent with a role for phytochromes in mediating a night-length signal, night-breaks are sufficient to restore flowering in SD grown plants (**Fig. S18**).

The results so far are consistent with phytochromes playing a role in perceiving night length and coordinating the photoperiod transcriptome via ELF3. Differential protein stability of phyC or PPD1/PRR37 would represent a potential mechanism to achieve photoperiodism. However we observe constantly levels of these proteins when they are overexpressed regardless of ZT time, suggesting modulating phyC protein levels is not key (**Fig. S19**). Since phytochromes in the active, Pfr, state slowly revert to the inactive Pr state in the dark (thermal or dark reversion), we hypothesized that this presents a mechanism for measuring the length of the night. Under long photoperiods the dark period may be insufficient for phyC Pfr to be depleted, with the result that ELF3 cannot accumulate to a high level. Extending the night period in short days however may enable phyC Pfr to become depleted, allowing the accumulation of repressive ELF3. To test this, we measured the dark reversion dynamics of Brachypodium phyC by overexpressing the gene in Brachypodium and Arabidopsis seedlings. In both cases, we observe similar reversion rates, and the dark reversion of phyC Pfr has a half-life of 8.3 h in Brachypodium (**Fig. 4H**; **Supplementary Dataset S6**). Under our long day conditions therefore, 72 % of Pfr remains at the end of the night, while in short day conditions only 37 % of Pfr is present at ZT0. This is consistent with the differences we observe in the binding of ELF3 to promoters in response to photoperiod, and suggests a model whereby the levels of Pfr provide a readout of night-length and communicate this to a central regulator of photoperiodism, ELF3 (**Fig. S20**).

## Discussion

Photoperiodism provides plants with important seasonal information to control their behaviour. As well as fundamental differences, such as long and short day plants, there is also considerable variation in the relative contribution of different factors to photoperiod sensitivity. Arabidopsis measures day-length to activate flowering (Hayama et al., 2017), and the phasing of the circadian rhythm with the external light-dark cycle is particularly important (Roden et al., 2002). In Brachypodium, wheat, rice and poplar, night-length is a critical determinant of photoperiodism (Ishikawa et al., 2005; Jos Ramos-Sánchez et al., 2019; Pearce et al., 2017). We are able to show that phyC plays a central role in this process, and the dark reversion rate of Pfr is consistent with a role for phyC as a “molecular hourglass” (Borthwick & Hendricks, 1960). While proposed about 60 years ago, this model was discounted on discovering a circadian variation in sensitivity to far red-light pulses during extended darkness (Cumming, Hendricks, & Borthwick, 1965). Our finding that phytochromes directly modulate the activity of the circadian component ELF3 suggests a mechanism to reconcile these observations. It will be interesting to see if the phyC-ELF3 signalling module is widely conserved to control photoperiodism, since this appears to be the case in rice (Itoh, Tanaka, & Izawa, 2018), and an ELF3 ortholog also controls photoperiodism in Pea (Rubenach et al., 2017).

Light quality at dusk varies seasonally (Hughes, Morgan, Lambton, Black, & Smith, 1984; Linkosalo & Lechowicz, 2006), and in Aspen phytochrome signaling controls growth cessation and budset during autumn (Olsen et al., 1997). The ability of phytochromes to integrate changes in both spectral quality and photoperiod as well as temperature (Jung et al., 2016; Legris et al., 2016) may represent a robust mechanism for making seasonal decisions.

## Acknowledgements

We thank Richard Amasino and Daniel Woods for sharing Brachypodium mutant lines. The work in KEJ’s laboratory is supported by the Leibniz Association. KEJ was supported by a fellowship from the Gatsby Foundation. PAW’s laboratory is supported by the Leibniz Association and the Leibniz IGZ.

---

**Fig. S1.**
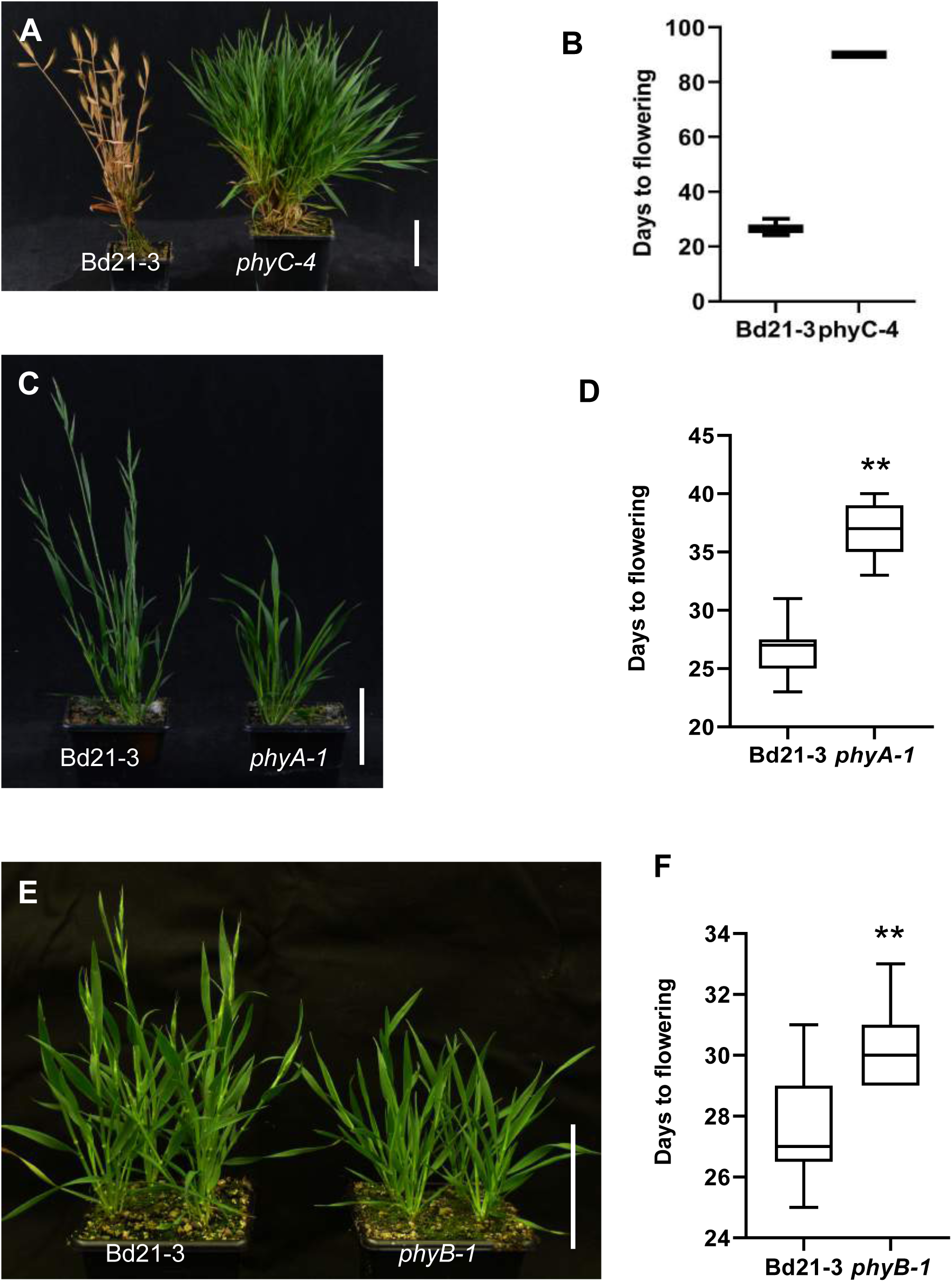
Phytochromes are necessary for LD activation of flowering. (**A** and **B**) *phyC-4* does not flower in inductive conditions. (**C** and **D**) *phyA-1* is late flowering in long days. (**E** and **F**) *phyB-1* is late flowering in long days.

**Fig S2.**
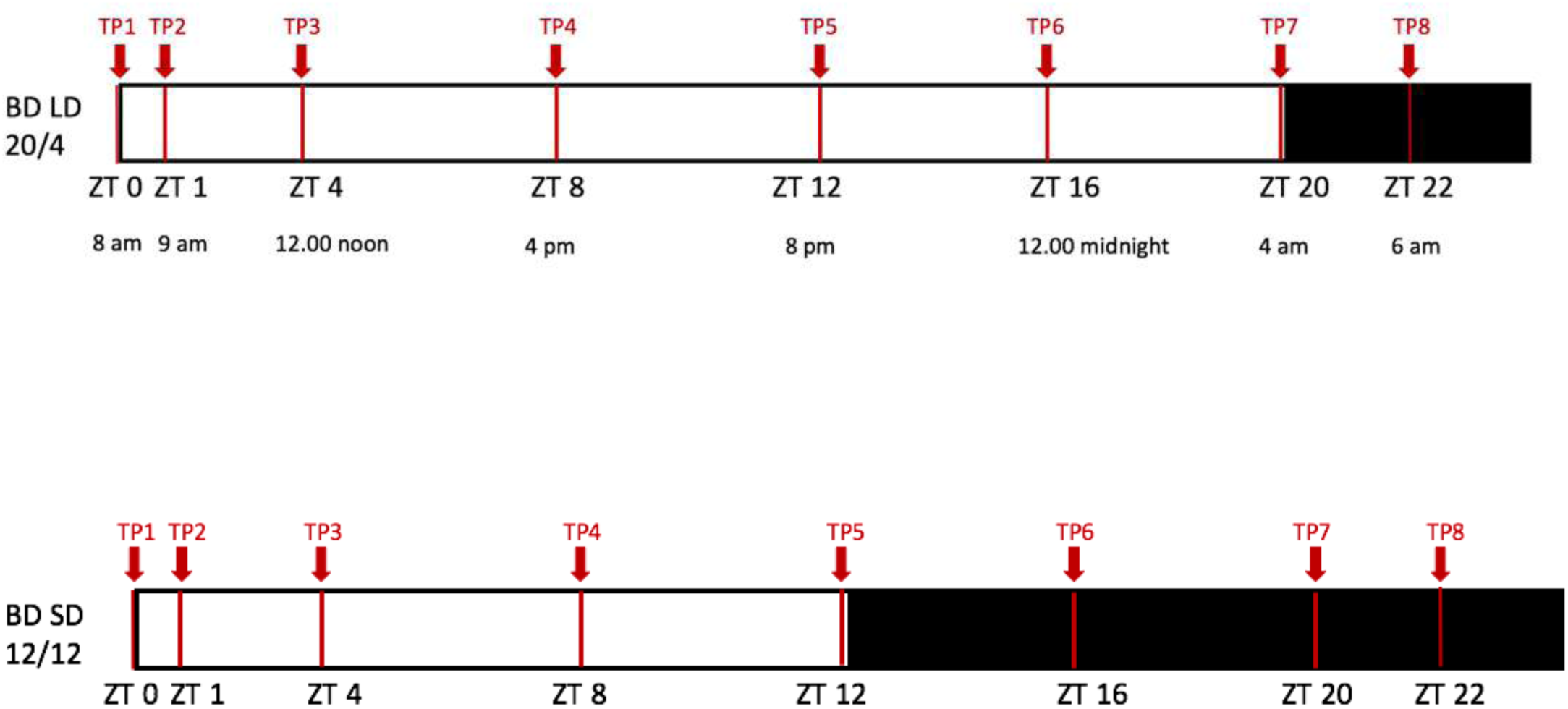
Sampling scheme for ChIP-seq and RNA-seq experiments. Schemata for collection points of all RNA-Seq, ChIP-Seq and Western blot series conducted in this study

**Fig. S3.**
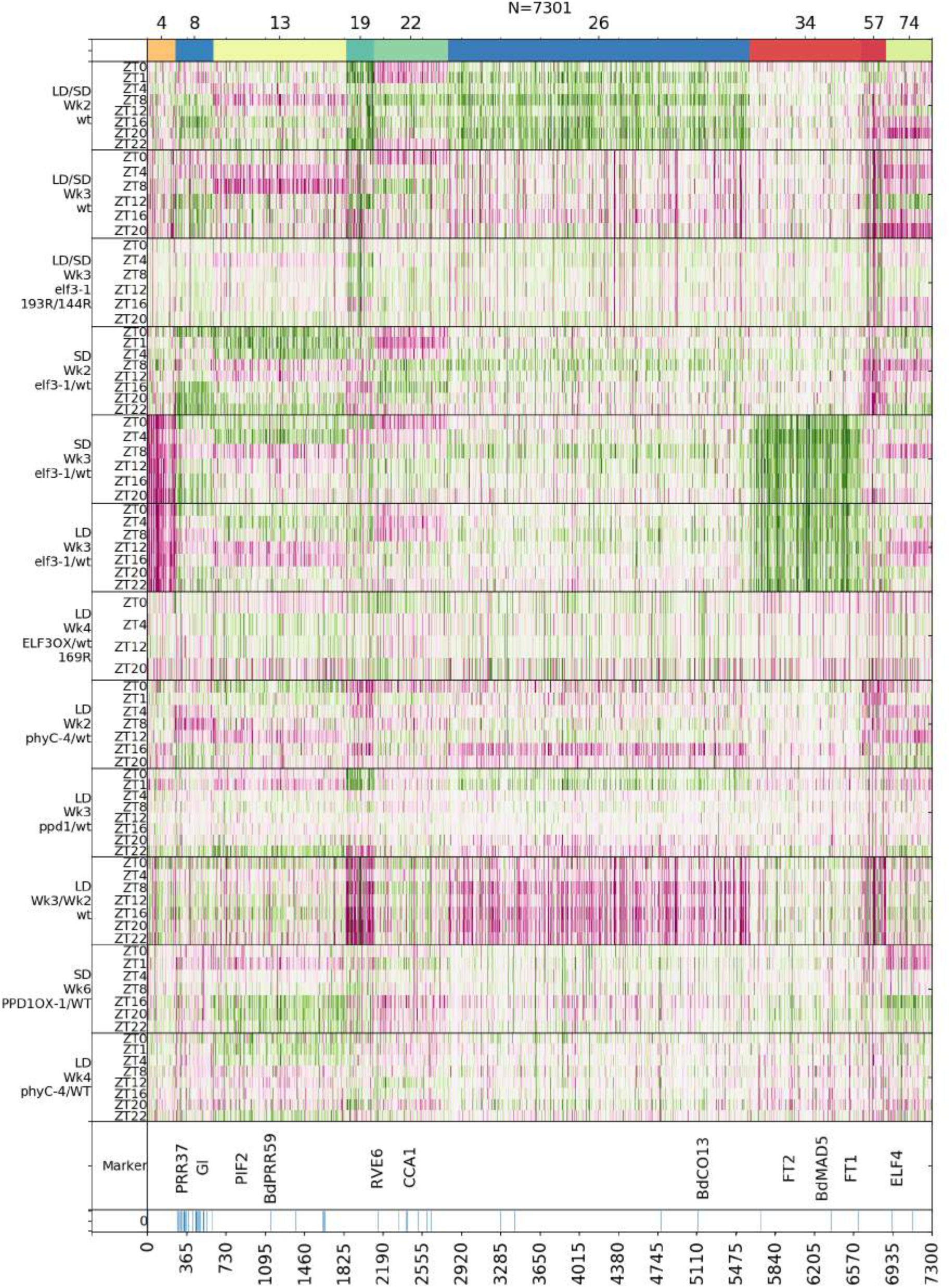
*PPD1* is necessary for the full induction of many clusters of gene expression that respond to photoperiod. Clustering of RNA-seq time-course data for *ppd1-1* plants grown in LD compared to WT. Multiple clusters including 4, 8, 13, 19, 22 and 26 show reduced expression in *ppd1-1* compared *to Bd21-3.* Genes which are upregulated are shown in green, genes which are downregulated are shown in red and unchanged is depicted in white.

**Fig. S4.**
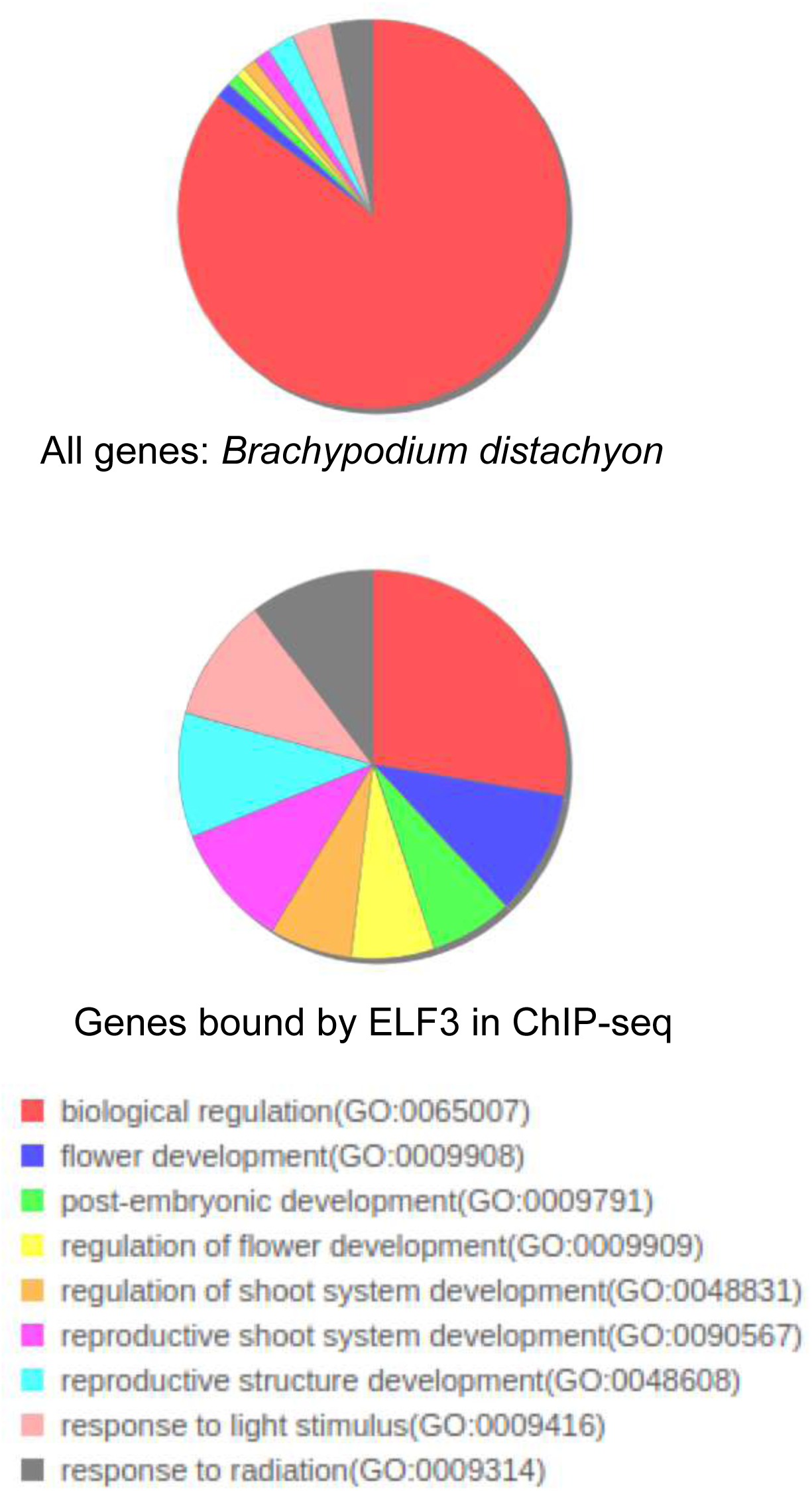
GO-enrichment for 34 genes bound by ELF3 and differentially expressed in *elf3-1*. We identified genes which are up-regulated significantly in elf3-1 compared to wild-type in SD in our transcriptome time-courses. Of these 1169 transcripts, we identified 34 that are directly bound by ELF3. This set of 34 genes that are both bound by ELF3 and whose transcription is regulated by ELF3 is enriched for genes involved in the regulation of flower development, reproductive shoot development as well as response to light, as has been observed in Arabidopsis (gene list in “Datasets”, DatasetS5).

**Fig. S5.**
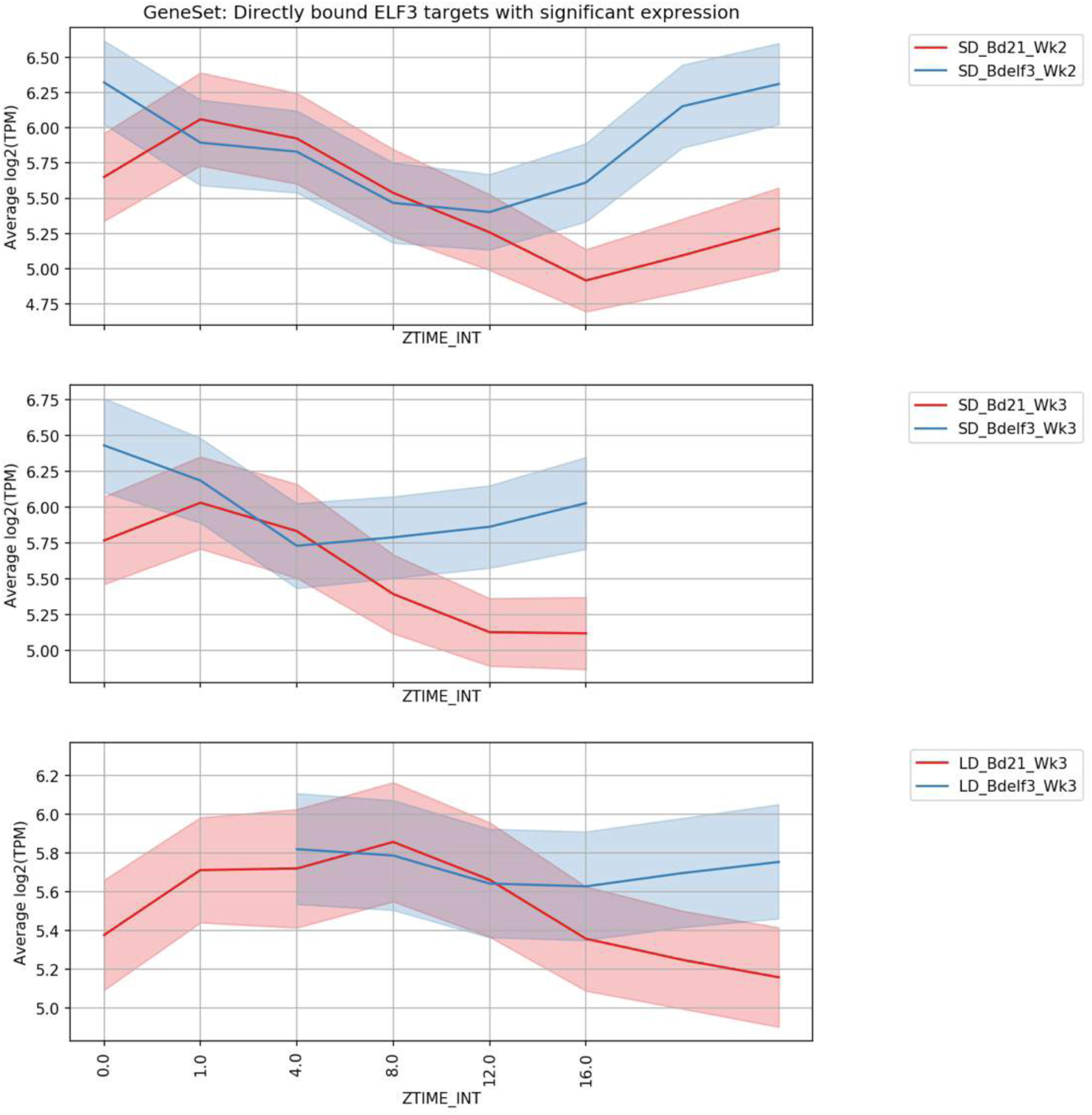
ELF3 is a transcriptional repressor. Genes bound by ELF3 show repression at the end of day and during the night. In the absence of ELF3 activity, these genes are no longer repressed during the night but are highly expressed. This indicates that the direct binding of ELF3 causes transcriptional repression. Y axis shows average expression in log scale and x-axis shows ZT, with start at ZT0

**Fig. S6.**
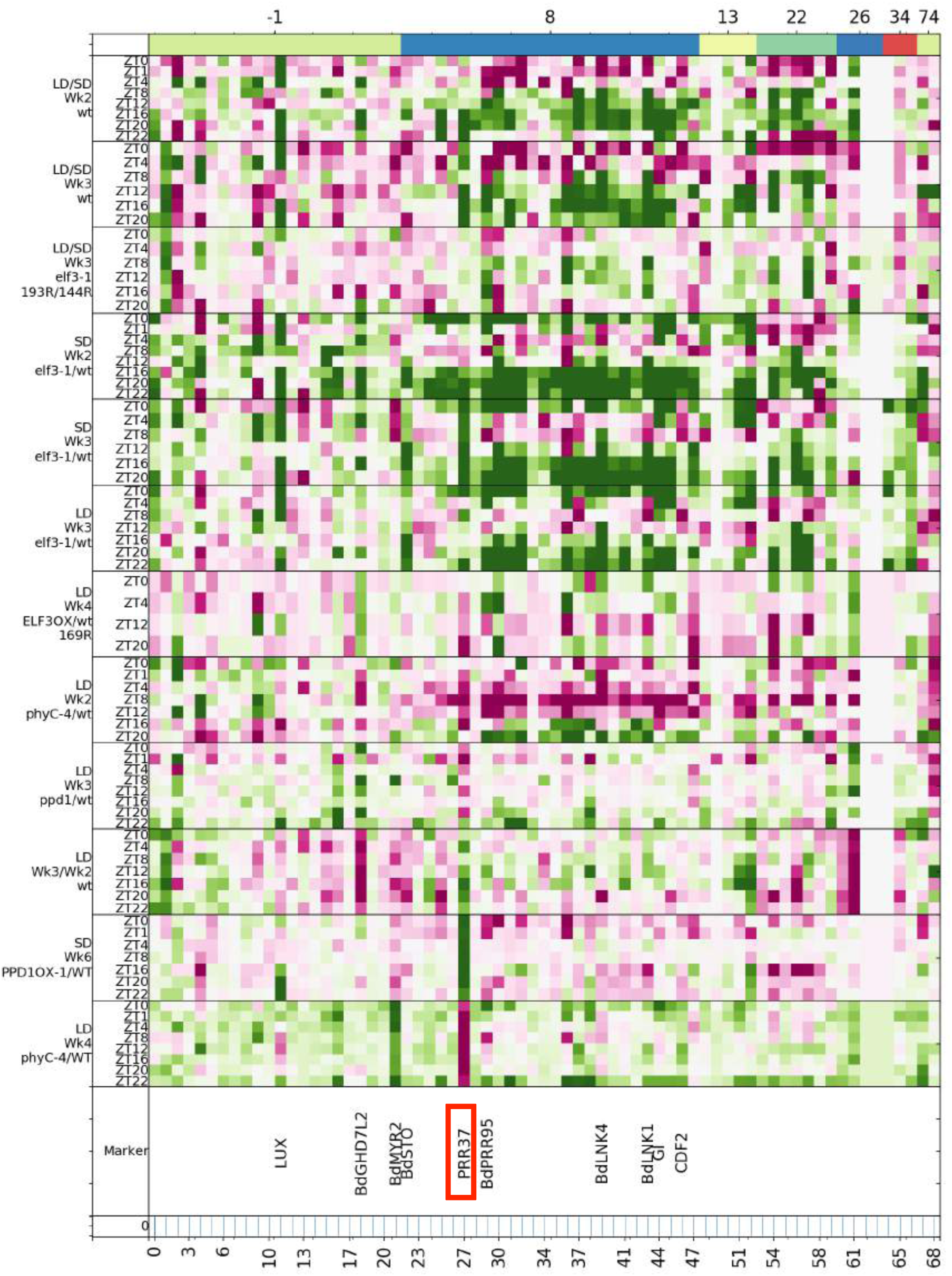
ELF3 bound genes show coordinated responses to photoperiod and genetic perturbation. Many of the genes that are directly bound by ELF3 are strongly induced by long photoperiods and in *elf3-1* (green). These genes are transcriptionally repressed in *phyC-4* in LD compared to WT in week 2. *PRR37/ PPD1* (red box) shows a particularly strong response to photoperiod, *elf3-1* and *phyC-4*. Perturbing *PRR37/PPD1* does not have as large a global effect on the transcriptome as for *ELF3* and *PHYC*, suggesting it is downstream in the pathway.

**Fig. S7.**
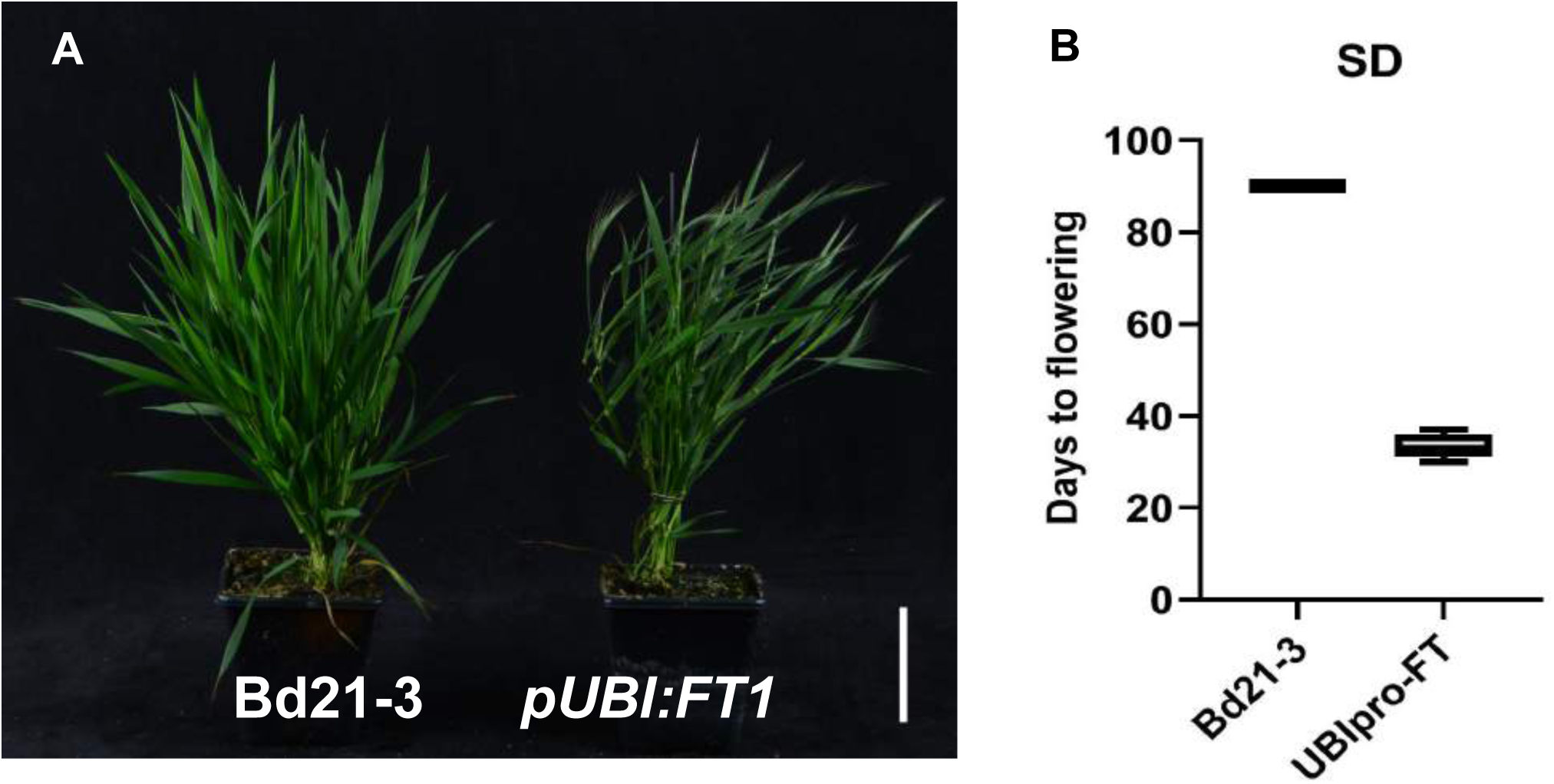
*Brachypodium* line overexpressing *FT1* (Bradi1g48830) under the UBI promoter (pUBI) flower almost independently of photoperiod. (*A) FT1-OX* under non-inductive SD conditions (12L:12D), Left: Bd21-3; Right: *pUBI:FT1*, Bar = 5 cm (B) *FT1-OX* plants flower nearly as early under non inductive short day conditions as under long days, whereas wild-type plants do not flower in 12L:12D.

**Fig. S8.**
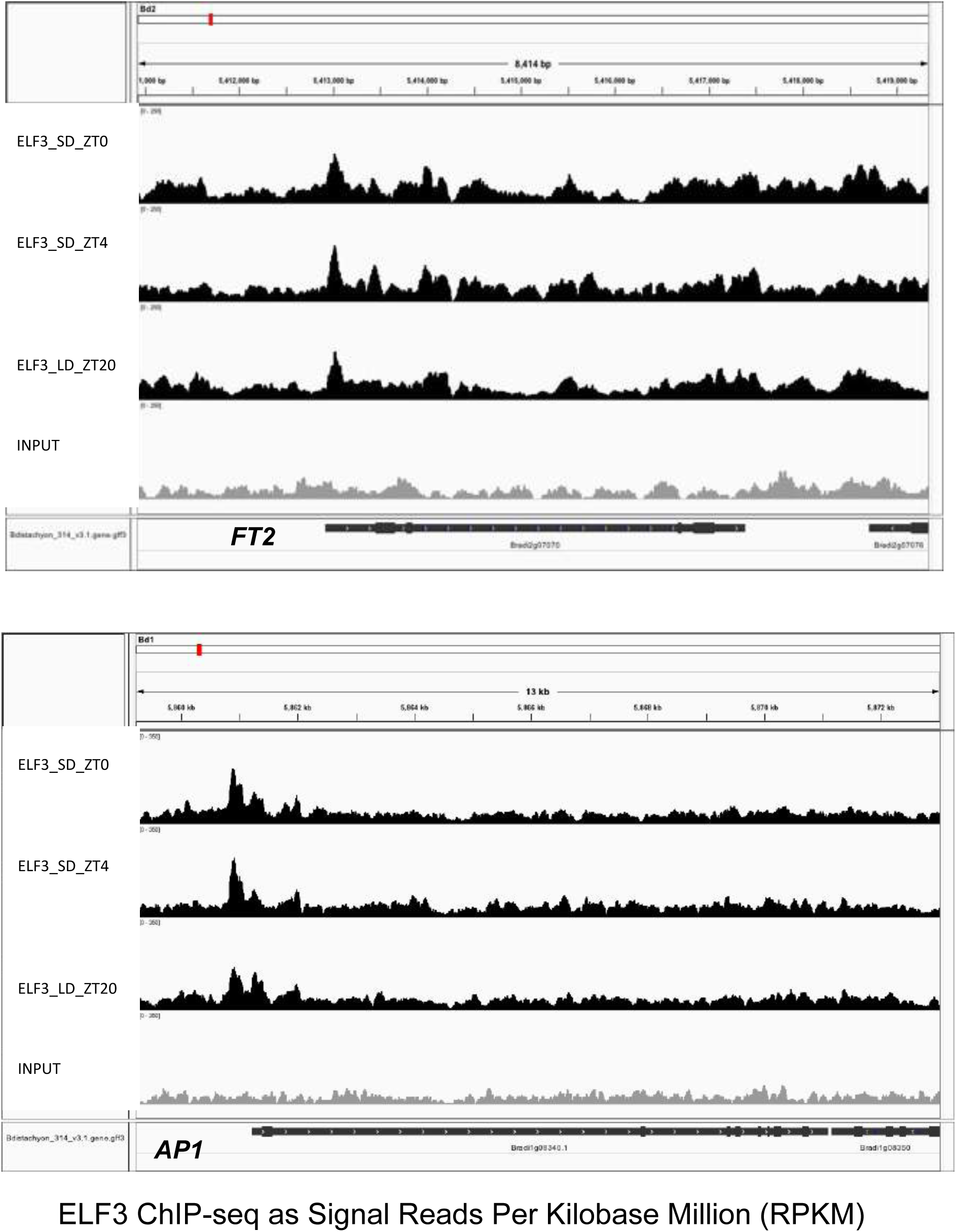
ELF3 ChIP-seq signal at key promoters. Shown are ELF3 ChIP-seq on the promoters of *FT2* and *AP1* at ZT0, ZT4 and ZT20 (SD, 12:12 and LD (20:4). 3 plants were used per sample. Last row shows the Input control. Height of tracks are shown as enrichment (Reads Per Kilobase Million, RPKM) in IGV browser.

**Fig. S9.**
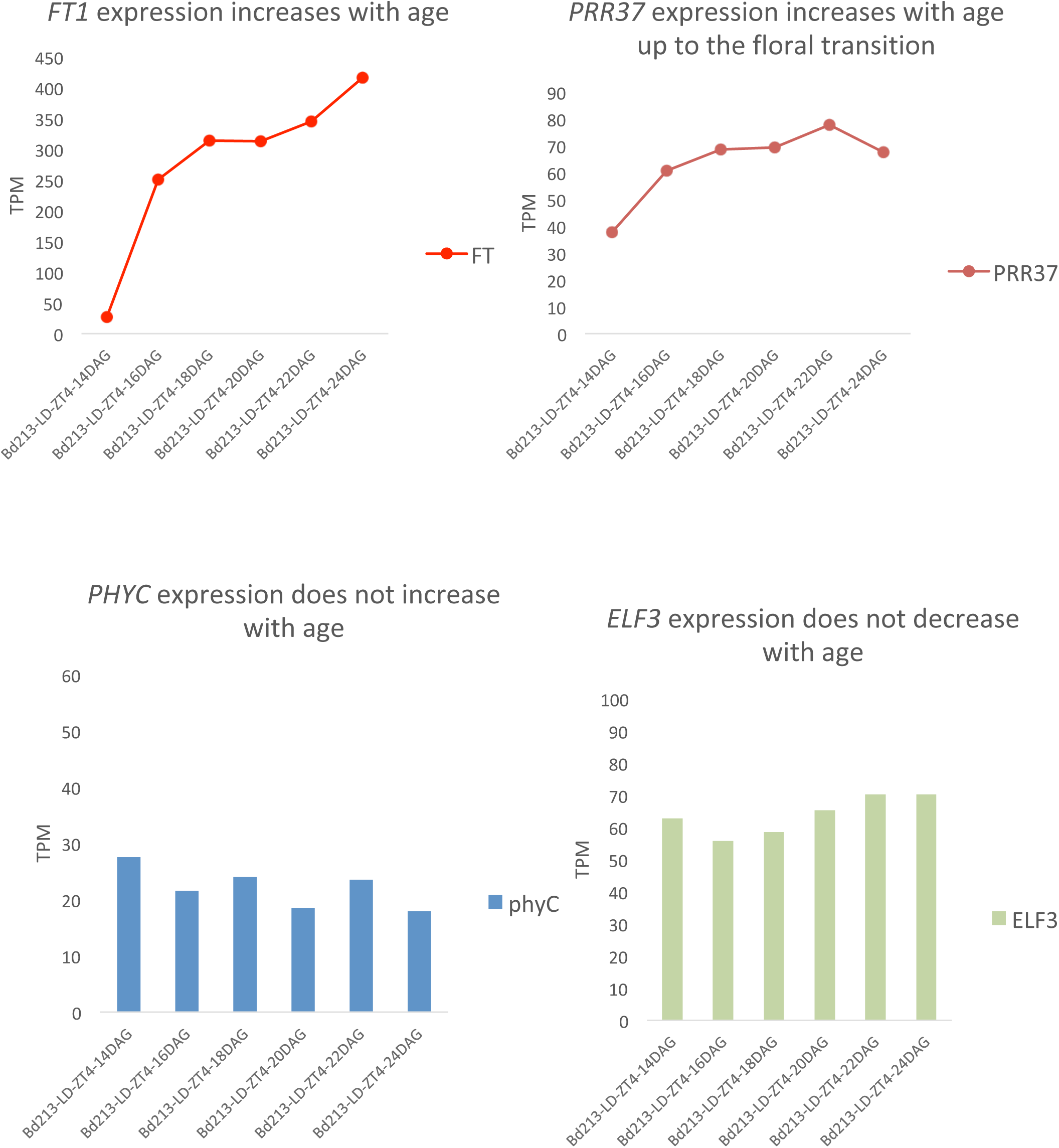
*FT1* and *PRR37* expression increase with age over a 14 day period. Plants were grown in LD and samples were collected every 2 days at ZT4 starting at 14 DAG, with 3 plants per sample. Expression is shown as transcripts per million (TPM). By comparison, *PHYC* and *ELF3* expression remains largely constant with age.

**Fig. S10.**
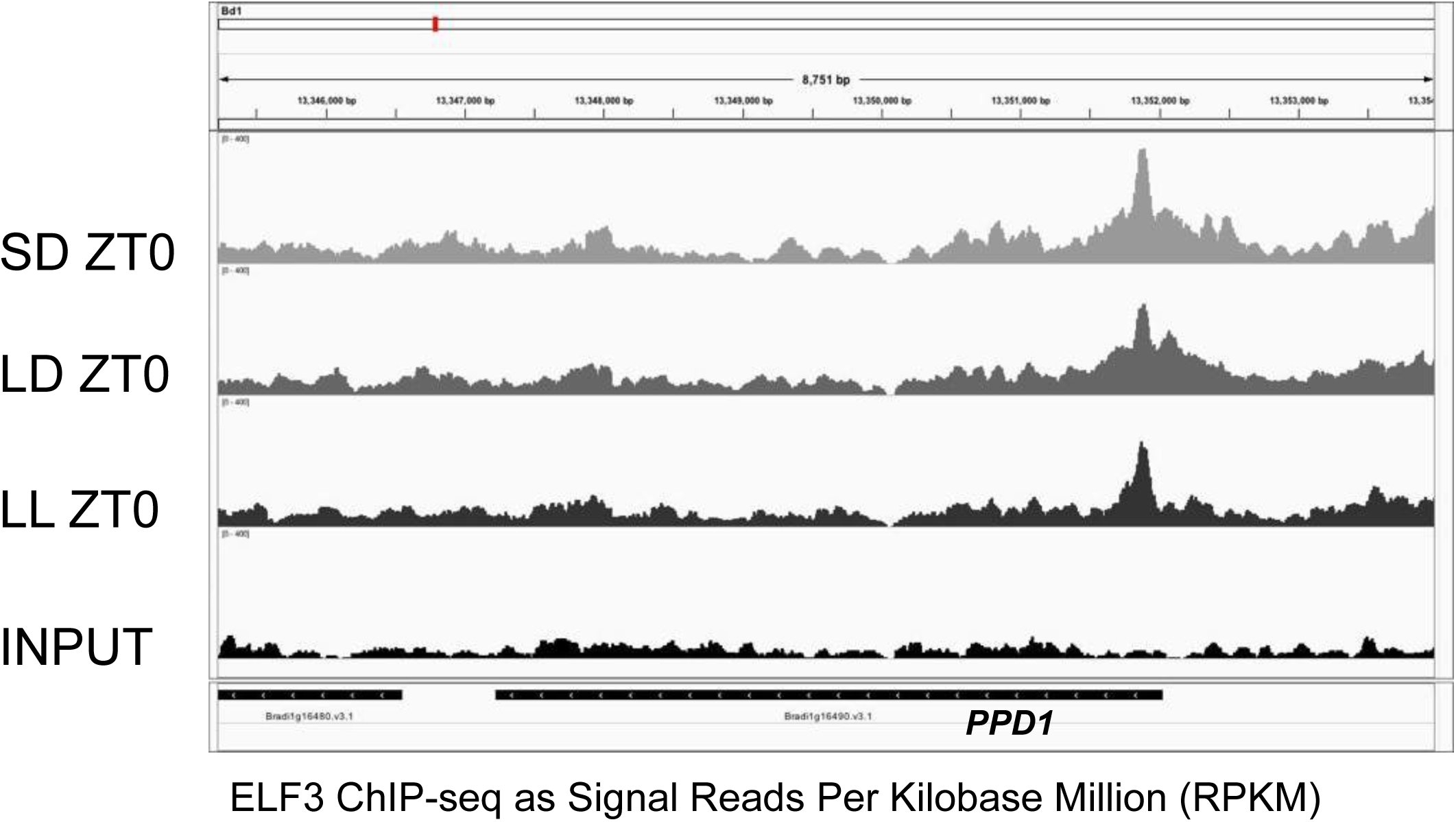
ELF3 binds at the PPD1 locus as assayed by ChIP-seq. Plants were grown (top to bottom) under SD, LD or LL conditions and sampled at ZT0 12 DAG, with 3 plants being used per ChIP-seq sample. Last row shows the Input control. Height of tracks are shown as enrichment (RPKM) in IGV browser. ELF3 binding is reduced in LL compared to SD. ELF3 binds PPD1 just upstream of the ATG. Analyses of *ppd1* alleles indicate that promoter insertions and deletions have played a major role modulating *PPD1* expression, revealing a 95 bp region within the promoter just upstream of the ATG that is conserved between wheat, barley and Brachypodium (Seki et al., 2013; Wilhelm, Turner, & Laurie, 2009), and it had been hypothesized that a photoperiod dependent repressor may bind this 95 bp region in short days to inhibit flowering.

**Fig. S11.**
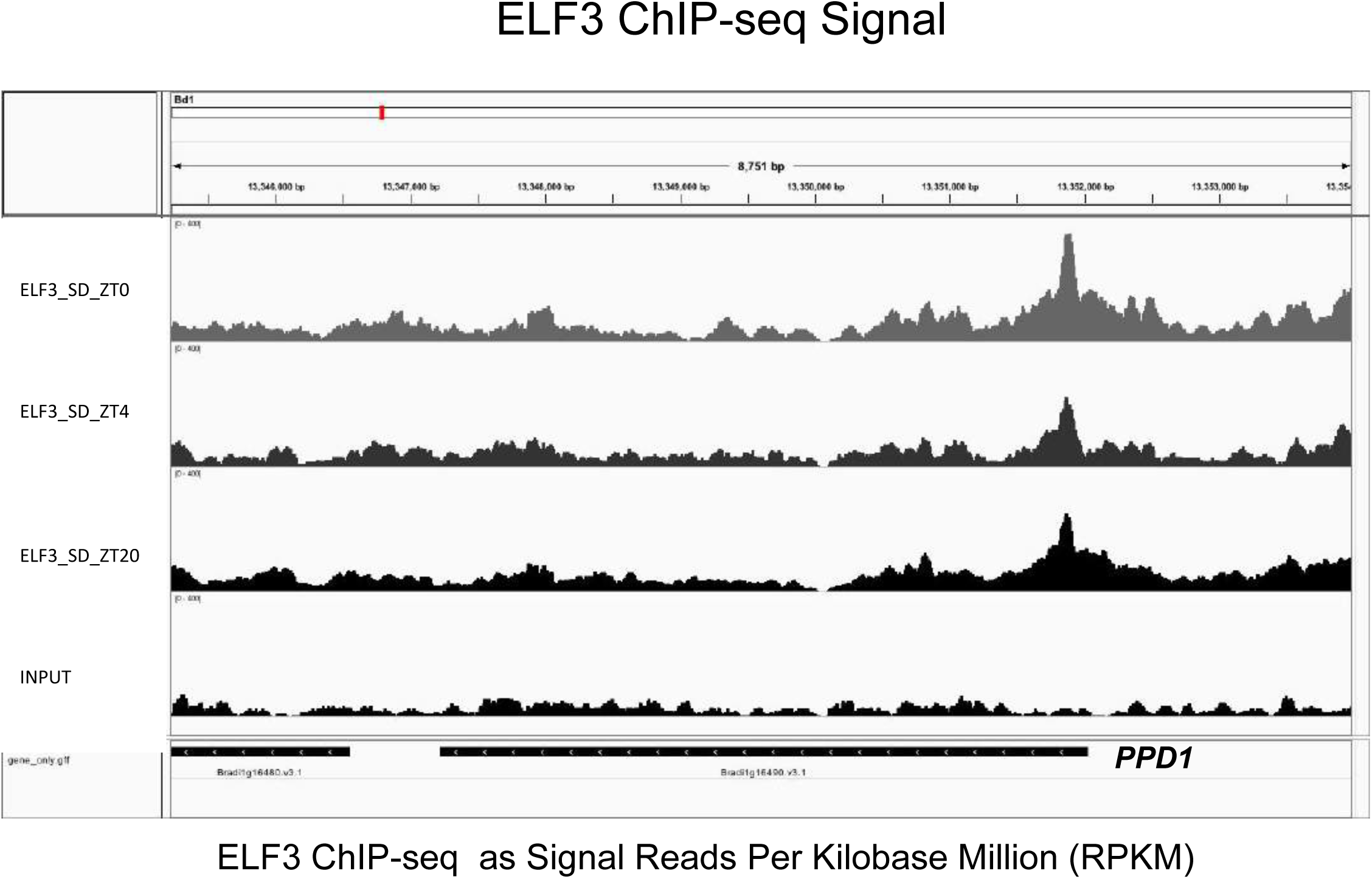
ELF3 binding at the *PPD1* locus assayed by ChIP-seq at ZT0, ZT4 and ZT20 in SD. Plants were grown in SD (12L:12D) and sampled 12 DAG, with 3 plants per sample. Last row shows Input control. ELF3 shows more binding during the night (ZT0 and ZT20) on *PPD1* promoter. Height of tracks are shown as Reads Per Kilobase Million (RPKM) in IGV browser.

**Fig. S12.**
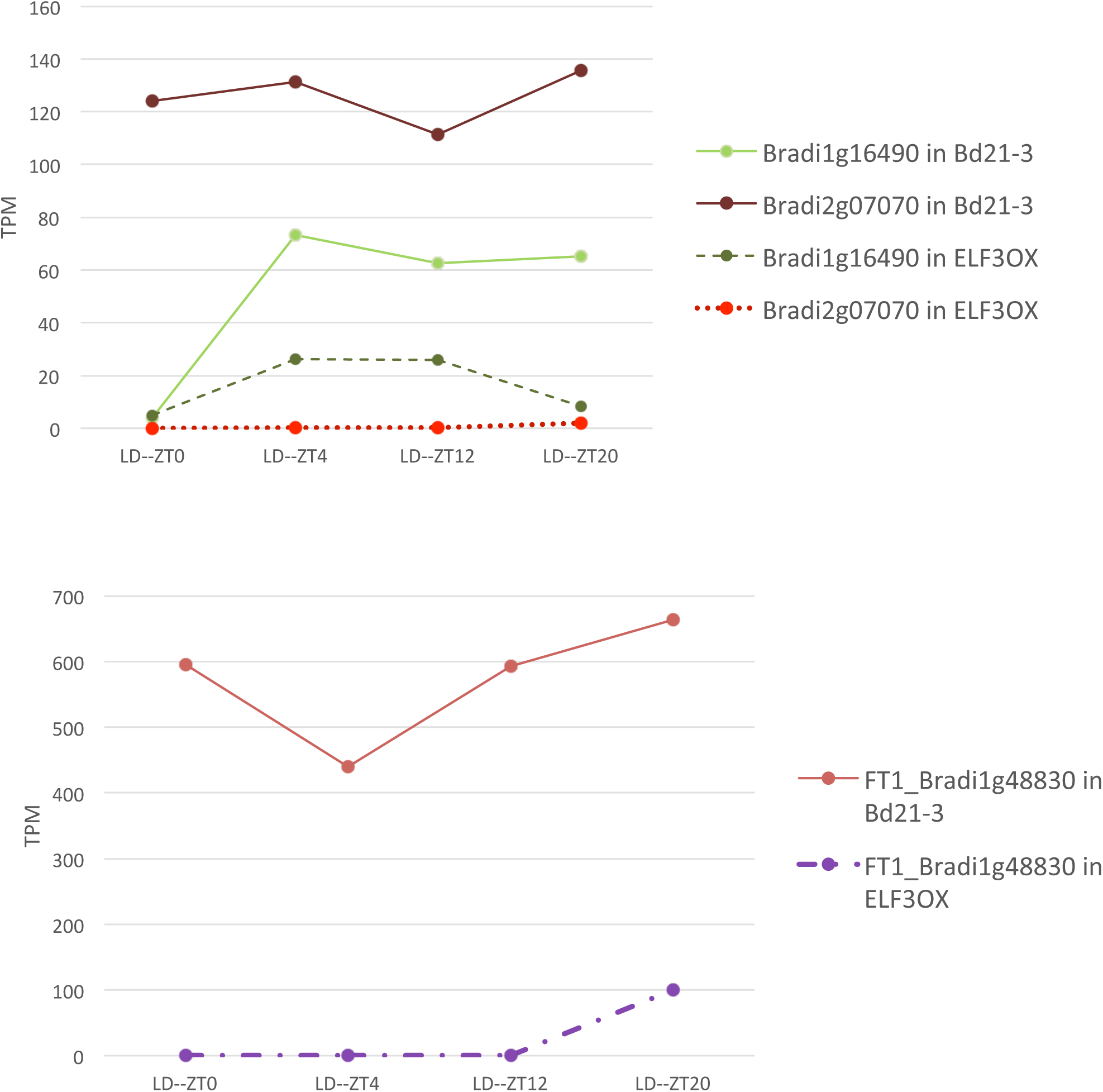
*ELF3* overexpression (*ELF3-OX*) represses downstream target gene expression. Expression levels of ELF3 bound genes (*PPD1*, Bradi1g16490; *FT2*, Bradi2g07070), and a downstream target (*FT1*, Bradi1g48830) are repressed in an *ELF3* overexpression line over a 24 h time course. Plants were grown under long day conditions and samples collected 3 weeks after germination at the indicated time. Values are given as transcript per million (TPM).

**Fig. S13.**
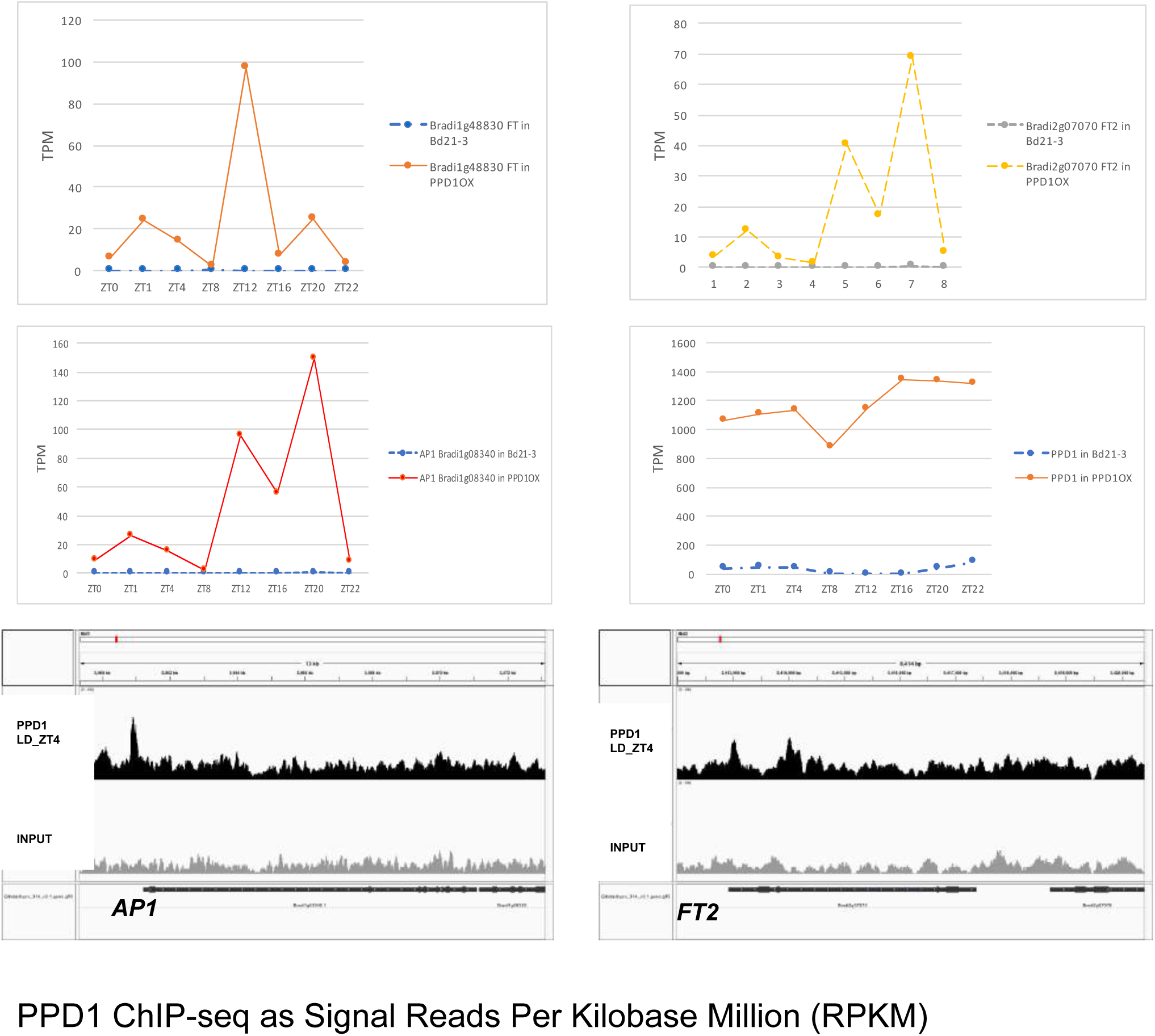
Expression levels of marker genes in the *pUBI:PPD1-Flag* line and ChIP-seq of PPD1. *FT1* (Bradi1g48830), *FT2* (Bradi2g07070) and *AP1* (Bradi1g08340) are highly upregulated in a *PPD1-OX* line under non-inductive SD conditions (12L:12D). Plants were grown for 6 weeks under SD condition and sampled at the indicated time, 3 leaves from individual plants were mixed for each sample. RNA-seq results shown as transcript per million (TPM). **PPD1 binds directly to 5’ region of *FT2* and *AP1*** as assayed by ChIP-seq. Top row (black) shows *PPD1-OX-Flag* ChIP-seq, and grey row shows Input control. Displayed in IGV browser.

**Fig. S14.**
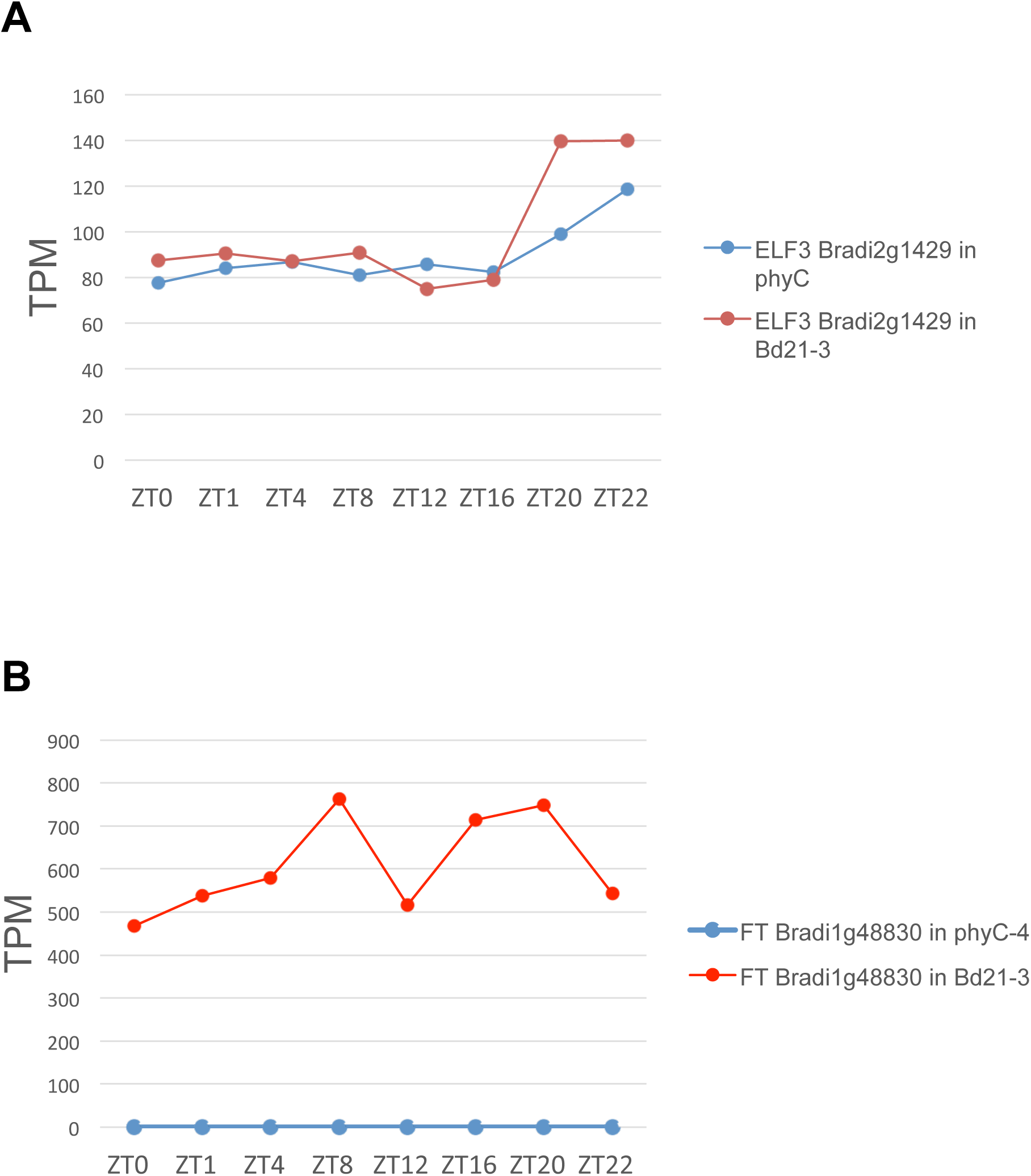
*ELF3* expression is not affected in *phyC-4, but FT1* expression is abolished. Plants were grown under long day conditions (20L:4D) and samples collected 4 weeks after germination at the indicated time. **A.** *ELF3* expression. **B.** *FT1* expression. Values are given as transcript per million (TPM).

**Fig. S15.**
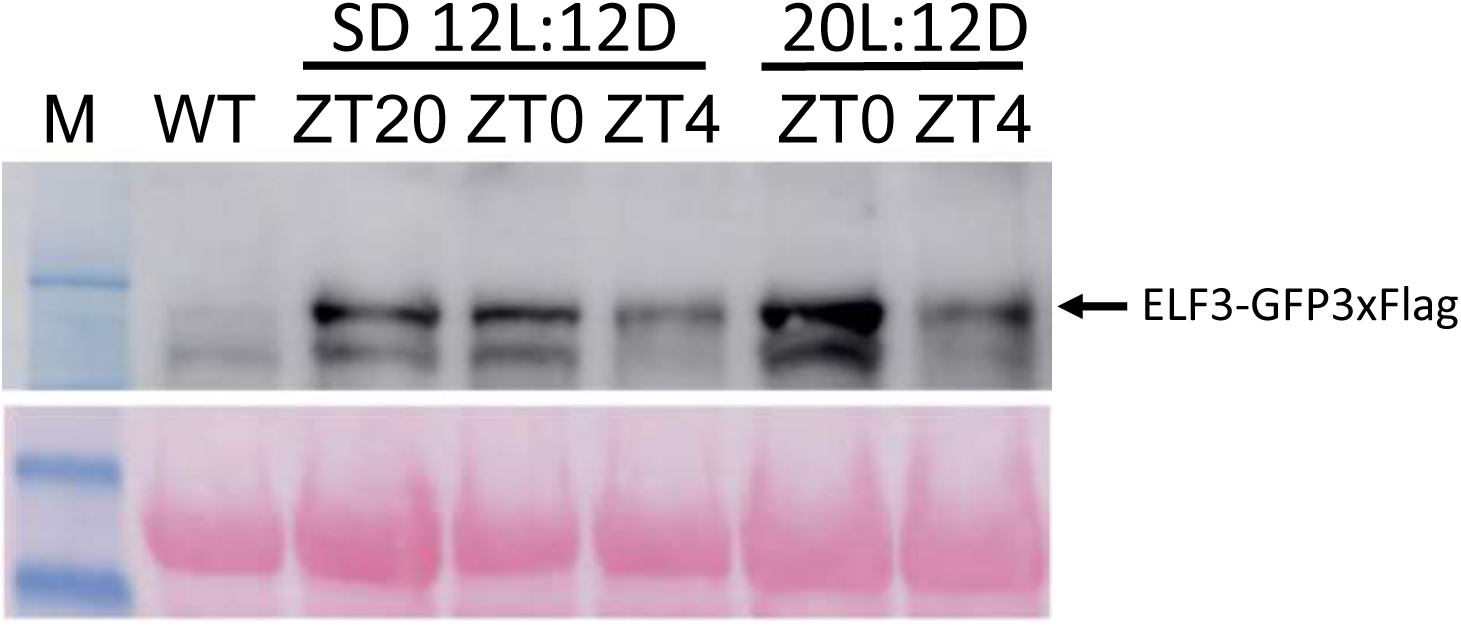
ELF3 protein is degraded in response to light. Additional Western blots showing degradation of ELF3 in response to light with antibody against ELF3 under SD (12:12) or 20L:12D condition. Plants were grown under 12L:12D (SD) or 20L:12D condition as indicated and samples taken 12 DAG at the indicated time (ZT20, ZT0 and ZT4, with 3 plants used per sample). We used wild-type plants (lane 2) or plants overexpressing ELF3 *(pUBI:ELF3-GFP-Flag)* (lane 3 to lane 7) and probed with an antibody raised in rabbit against ELF3 peptide *(Agrisera)*.

**Fig. S16.**
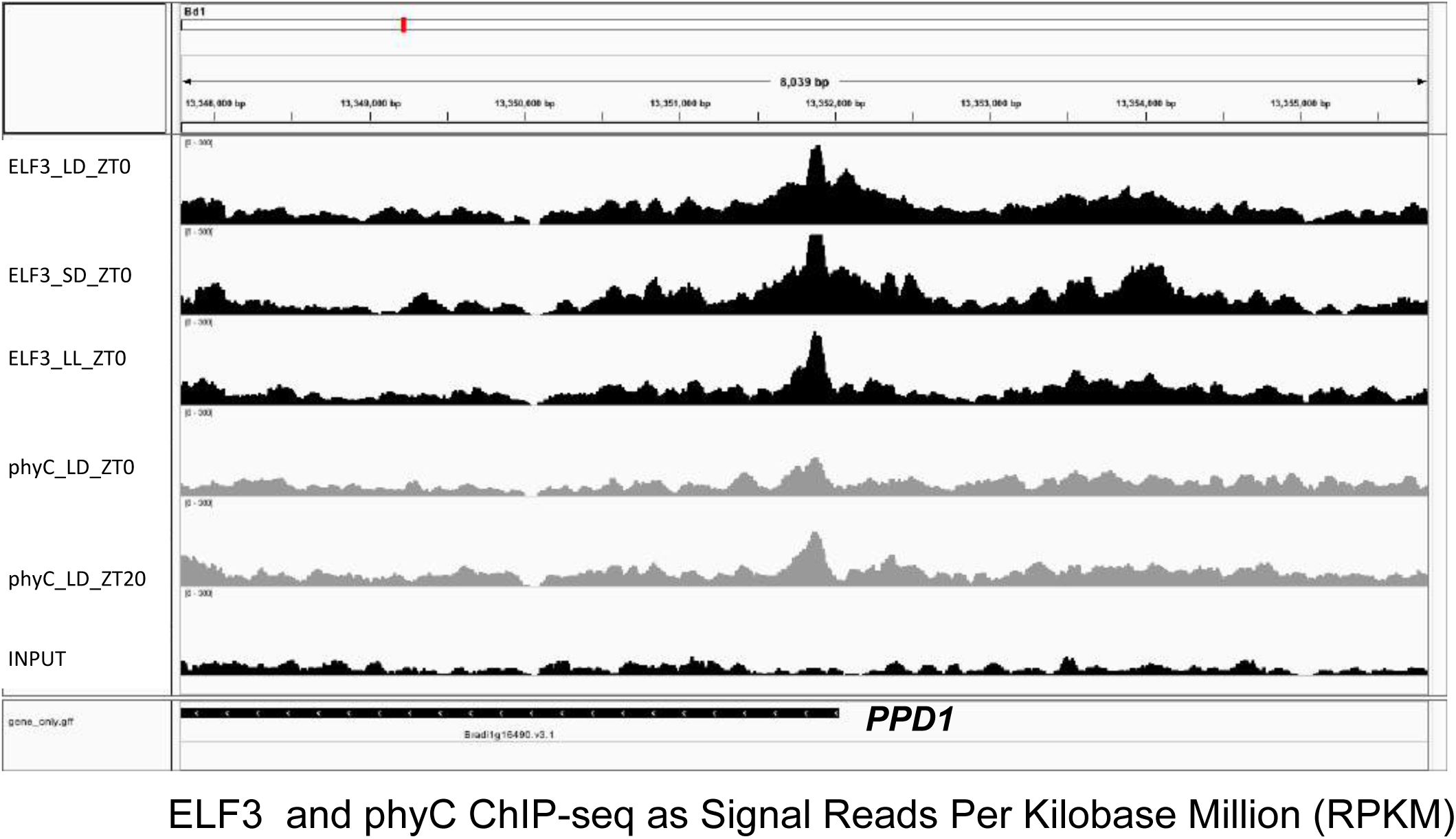
Enrichment of ELF3 and phyC at the *PPD1* promoter. Plants were grown (top to bottom) under LD, SD or LL conditions and sampled at ZT0 12 DAG (ELF3) or ZT0 and ZT20, LD (phyC), with 3 plants being used per ChIP sample. Last row shows the INPUT control. Height of tracks are shown as enrichment (RPKM) in IGV browser.

**Fig. S17.**
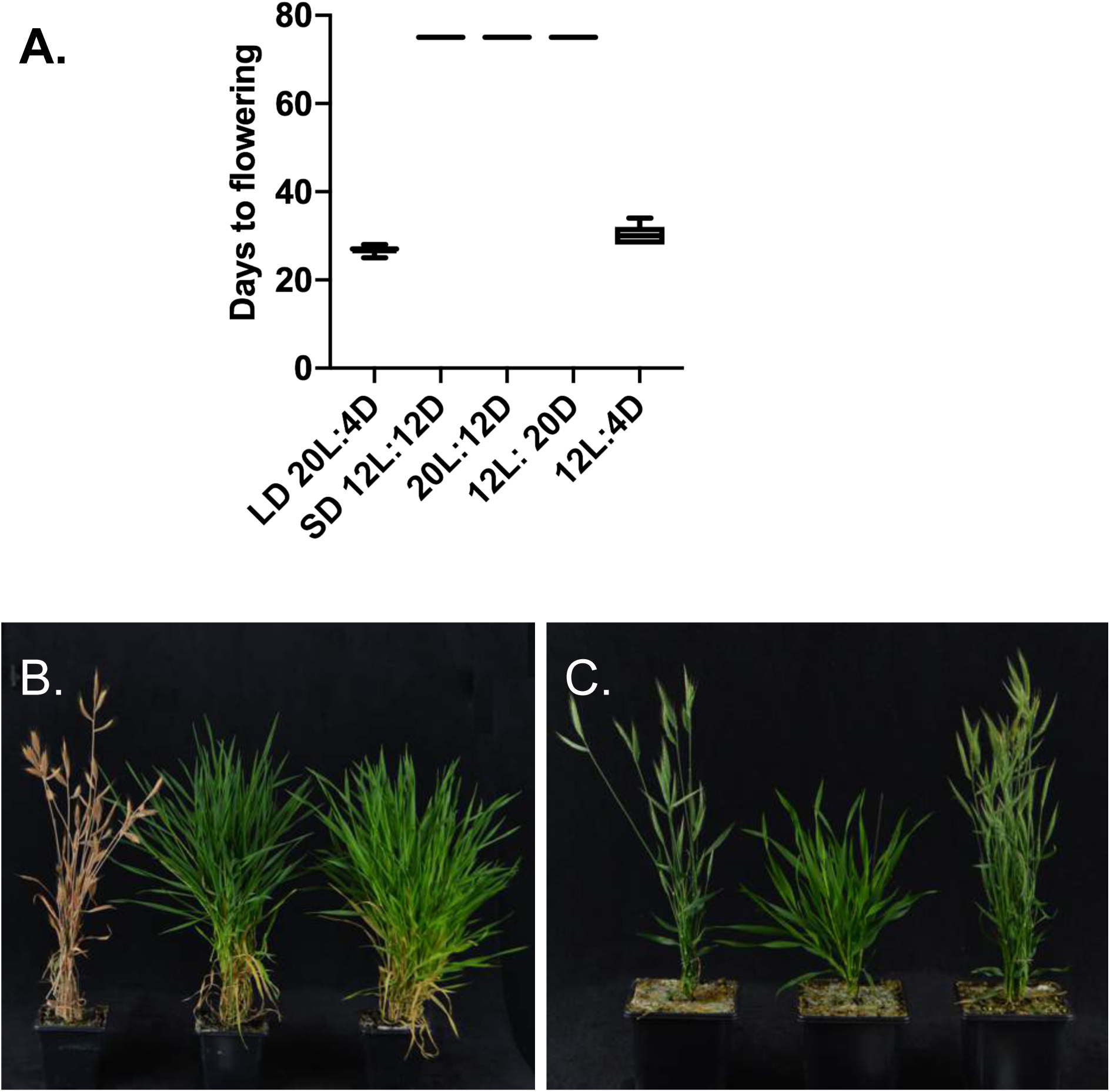
Night length, but not day length, determines flowering in *Brachypodium.* **A.** Plants grown under 12 h light and 4 h dark (12L: 4D) condition flower nearly at the same time as plants grown in 20L:4D (LD) conditions, whereas plants grown under 12L:12D, 20L:12D or 12L: 20D did not flower during the course of the experiment. Experiment was terminated at 75 days, when plants began to senesce. *Brachypodium* does not flower when grown under long night conditions, regardless of day length. **B.** From left to right: Bd21 grown under 20L:4D (LD), 12L: 20D and 20L:12D. **C.** Brachypodium flowers early when grown under short night conditions, regardless of day length. From the left to right: *Bd21* grown under 20L:4D (LD), 12L:12D (SD) or 12L:4D

**Fig. S18.**
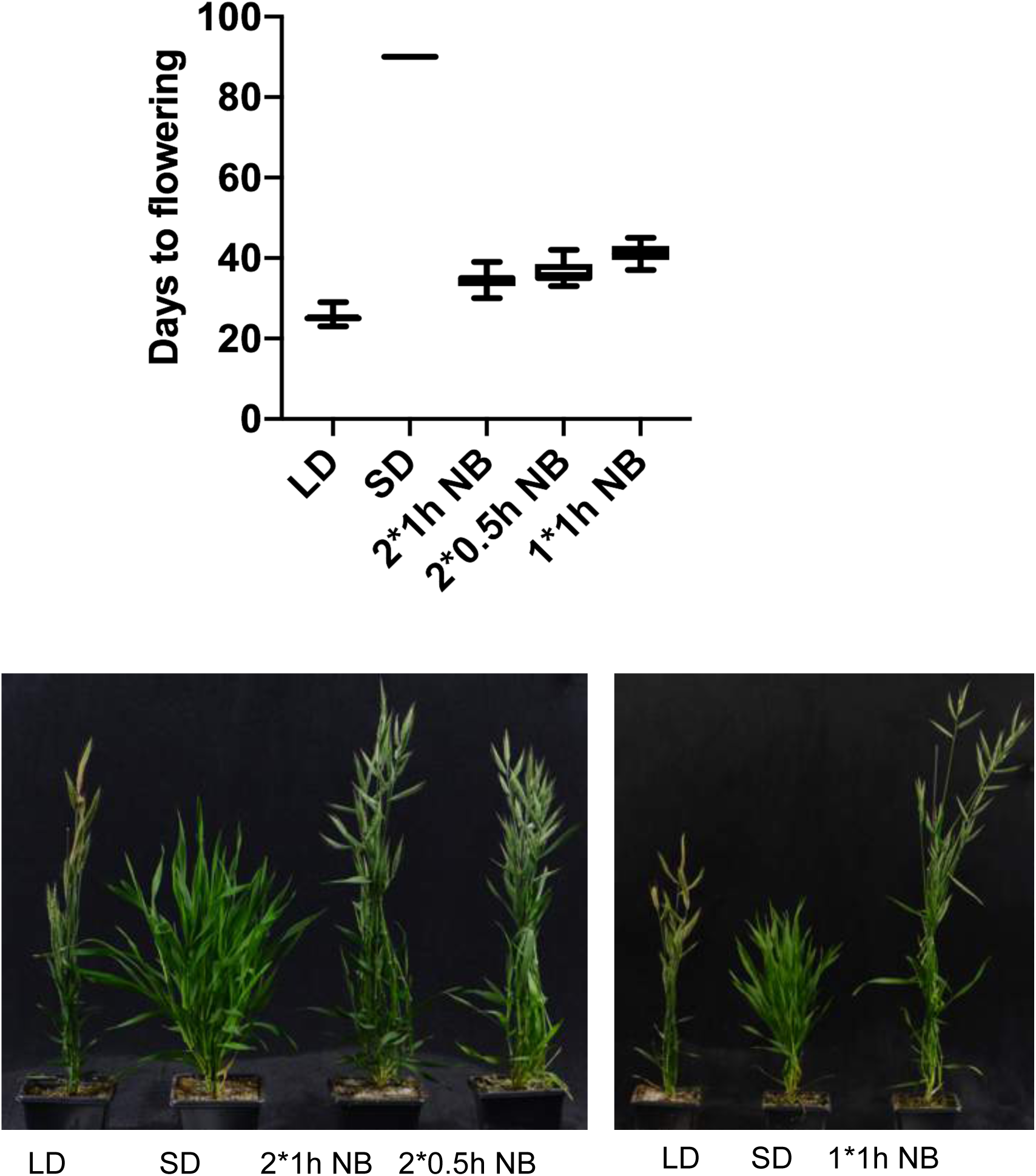
A night break is sufficient to trigger flowering under non-inductive short day conditions in *Brachypodium.* Introducing a single 1 h night break or 2 night breaks of 30 min results in a flowering time similar to a plant being grown under inductive long days. Conditions: **2*1h NB:** 12 hour light + 4 hours dark + 1hour light + 3 hours dark + 1hour light +3 hours dark. **2*0.5h NB:** 12 hour light + 4 hours dark + 0.5 hour light + 3.5 hours dark + 0.5 hour light + 3.5 hours dark. **1*1h NB:** 12 hour light + 6 hours dark + 1hour light + 5 hours dark, all Bd21

**Fig. S19.**
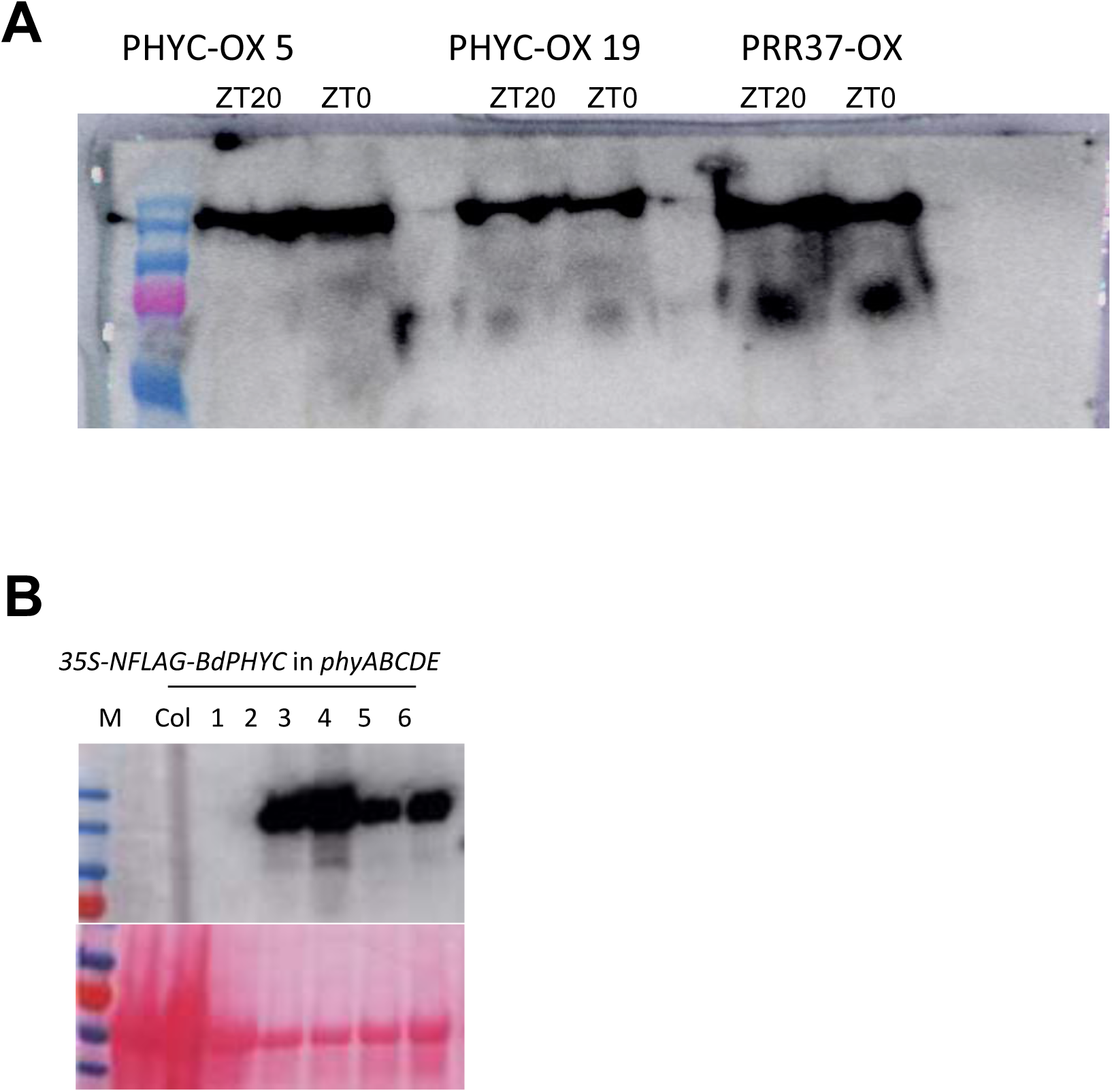
Detection of key proteins by immunoblot in plant extracts. **A.** phyC and PRR37 protein levels are stable and do not change at different times of day. Plants were grown under 20L:4D (LD) condition and samples taken 12 DAG at the indicated time (ZT20, ZT0 with 3 plants used per sample). We used 2 independent lines expressing *pUBI:phyC-GFP-Flag* and 1 expressing *pUBI:PRR37-Flag* probed with an antibody against Flag epitope (M2, Sigma). **B.** *35S-NFlag-BdPHYC* is stably expressed in *phyABCDE Arabidopsis.* We tested 6 independent lines for expression level of BdphyC. Western blot was probed with with an antibody against Flag epitope (M2, Sigma).

**Fig S20.**
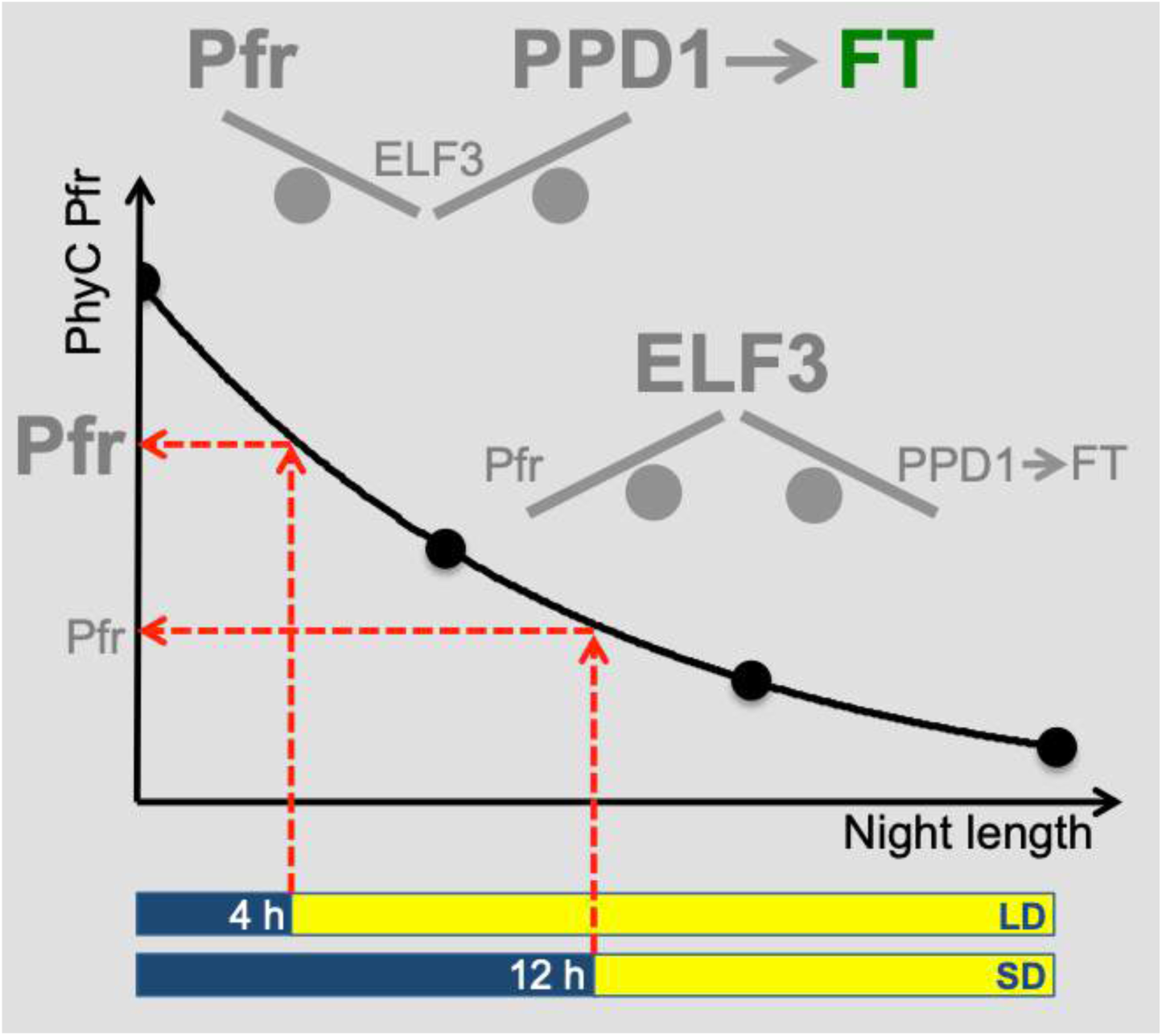
The dark reversion of phyC Pfr is suitable to provide a measure of the length of the night. During SD, PhyC Pfr declines to low levels during the longer nights, enabling greater accumulation of the floral repressor ELF3. In LD, nights are short and phyC Pfr levels remain high. This prevents ELF3 accumulation, allowing higher levels of *PPD1* to be expressed, allowing the activation of *FT*.

## Materials and Methods

### Plant materials and growth conditions

*Brachypodium distachyon* accession *Bd21-3* and *Bd21* were used in this study. Seeds were imbibed in distilled water at 4 ° Cfor two days before sowing. Plants were grown in 5 parts John Innes #2, 3 parts peat, 1 parts silver sand, 3 parts course vermiculite, Osmocote 2.7 g/L. All plants were grown in growth cabinets with constant temperature 20 °C, 65 % humidity and 350 µmol m^−2^ s^−1^ PPFD (Photosynthetic Photon Flux Density). For flowering-time experiments, plants were grown in photoperiod regimes: (a) LD (20 h light/4 h dark); (b) SD (12 h light/12 h dark); (c) 20:12 (20 h light/12 h dark); (d) 12:20 (12h light /20 h dark), (e) 12:4 (12 h light/4 h dark).

### Mutants used in this study

| Line Name | Background | Description | Source | Notes |
| --- | --- | --- | --- | --- |
| *elf3-1* | Bd21-3 | crispr line | This study | 7bp deletion or 1bp insertion in the second exon, both caused premature stop codon |
| *phyC-4* | Bd21-3 | crispr line | This study | 4bp deletion in the first exon, caused premature stop codon |
| *ppd1-1* | Bd21-3 | crispr line | This study | 1bp deletion in the sixth exon, caused premature stop codon |
| *phyA-1* | Bd21-3 | crispr line | This study | 3bp deletion in the first exon, caused one amino acid deletion |
| *phyB-1* | Bd21-3 | T-DNA | https://jgi.doe.gov/our-science/science-programs/plant-genomics/brachypodium/brachypodium-t-dna-collection/ | T-DNA insertion in the 5'UTR. |
| *UBIpro-ELF3* | Bd21-3 | transgenic line | This study |  |
| *UBIpro-PPD1* | Bd21-3 | transgenic line | This study |  |

The *phyC-1 EMS* mutant has been described previously (*1*). The *Arabidopsis thaliana* phytochrome mutants in Ler backgrounds *phyabde*, and *phyabcde* were provided by K. Franklin.

For this study we created CRISPR mutation in the *ELF3* gene (Bradi2g14290), *PHYC* gene (Bradi1g08400) and *PPD1* gene (Bradi1g16490). The cloning of the single guide RNA (sgRNA) was done as described in (*2*). sgRNAs primers for *ELF3*, *PPD1* and *PHYC* were designed using design tool http://www.e-crisp.org/E-CRISP/. The annealed oligos were ligated into entry vector pOs-sgRNA and then cloned into destination vector pH-Ubi-cas9-7 by gateway LR reaction. The constructs were transformed in the *Agrobacterium* strain AGL1. *Agrobacterium*-mediated plant transformation of embryonic callus generated from immature embryos was performed as described (*3*). For the genotyping analysis, mutations were confirmed by sequencing and T2 lines with mutation but not carrying Hyg resistance and were selected for further analysis.

For the overexpressing transgenic lines, the genomic coding sequence of *ELF3*, *PPD1* and *PHYC* were amplified by PCR with primers indicated in Table S1. The PCR products were cloned into SLIC binary vector containing Ubiquitin promoter and C-terminal 3xFLAG tag using NEBuilder® HiFi DNA Assembly Cloning Kit (E2621L). pENTR-YFP-His_6_-3xFLAG (*4*) was recombined using the Gateway system (Invitrogen) into pMDC32 (*5*). Embryogenic calli from *B. distachyon* 21-3 plants were transformed with pENTR-YFP-His_6_-3xFLAG as described (*6*). For each construct, approximately 20 independent transgenic lines were obtained and homozygous single insertion lines were selected for further analysis.

For overexpression of *PHYC* in Arabidopsis, the *PHYC* genomic fragment was amplified and then cloned into 35S and N-terminal 3xFLAG tagged binary vector by NEBuilder® HiFi DNA Assembly Cloning Kit (E2621L). The binary construct was transformed into Arabidopsis *phyABCDE* mutant by floral dipping method. The *35S-N3FLAG-PHYC* transgenic plants were isolated by Kanamycin selection and propagated to obtain homozygous seeds to measure the dark reversion rate.

**Table S1.**
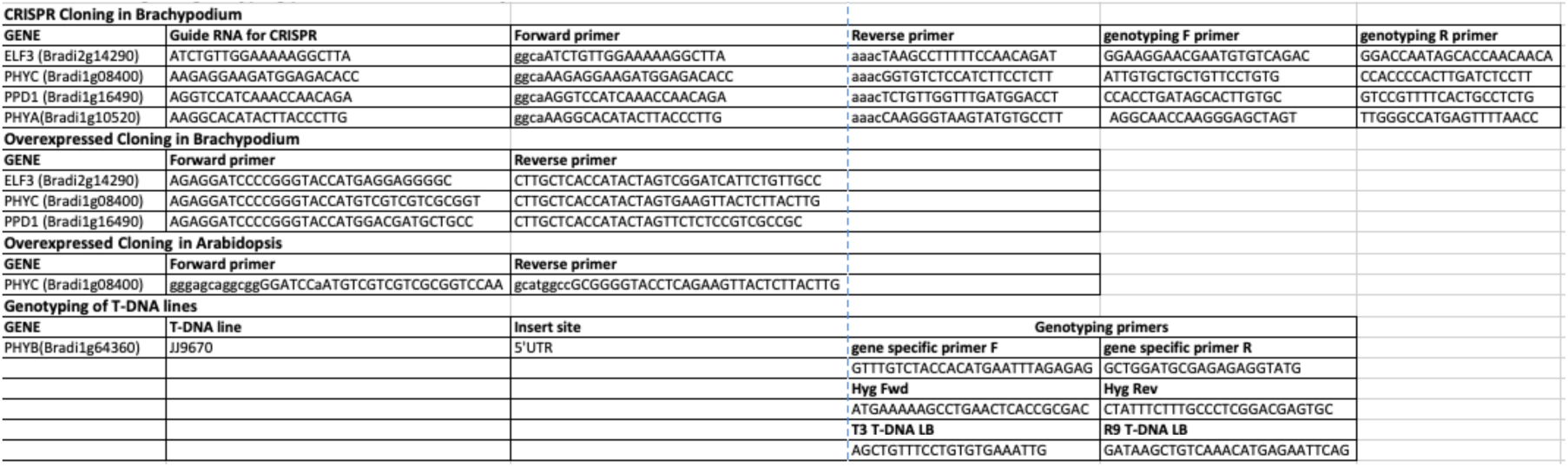
Cloning and genotyping primers used in this study.

For **Western blot assay**, seeds were sterilized and sown on ½ X Murashige and Skoog-agar (MS-agar) plates at pH 5.7 and grown in Magenta™ GA-7 Plant Culture Box (Thomas scientific). Sterilized seeds were stratified for 2 days at 4°C in the dark and allowed to germinate. The plates were transferred to short-day conditions (12 h light and 12 h dark) and collected at the indicated time.

3 plants per sample were grinded and 100 mg of frozen plant material then added 100 µl 2x Laemmli buffer (S3401, SIGMA). The protein was denatured at 96 °C for 10 min. 15 µl of protein samples were separated on 12 % SDS-PAGE and blotted 7 min to nitrocellulose membrane using Turbo semi-dry transfer. Blots were blocked with 5 % milk for 1 h at RT and then incubated in the anti-FLAG M2 (Sigma) primary antibody at a dilution of 1:2500 at 4 °C overnight with agitation. The antibody solution was decanted and the blot was rinsed briefly twice, then washed once for 15 min and 3 times for 5 min in TBS-T at RT with agitation. Blot was incubated in secondary antibody goat anti-mouse IgG-HRP conjugate (Bio-Rad, #1721011) diluted to 1:5000 in for 2h at RT with agitation. The blot was washed as above and developed by PiecreTM ECL substrate (Thermo Scientific, #32134). Exposure time was 15 and 30 seconds.

### Gene expression by RNA-seq (Fig. 2)

RNA-seq experiments were performed for Bd21-3, *elf3-*1, *UBIpro:ELF3* and *phyC-4*, *ppd-1*, *UBIpro:PPD1* at LD and SD over a 24h time course. 2, 3 or 6 week old seedlings of the indicated genotypes were grown at 20 °C and sampled at intervals over the diurnal cycle: ZT = 0, 4, 8, 12, 16, 20 and 22 h. For each sample 3 leaves of 3 plants were mixed. The *phyC-4* time course was done as replicate. In each time course performed a wildtype control included in all conditions and ZTs.

Qiagen RNeasy Mini Kit (74104) was used to extract RNA. RNA quality and integrity were assessed on the Agilent 2200 TapeStation system. Library preparation was performed with 1µg total RNA using the NEBNext® Ultra™ Directional RNA Library Prep Kit for Illumina® (E7420L). The libraries were sequenced on a NextSeq500 (Illumina) running a final pooled library. Each pool contained 24 to 30 samples and was sequenced using NextSeq® 500/550 High Output Kit v2 (150 cycles) TG-160-2002 on a NextSeq500 (Illumina).

**Q-PCR** was performed on a Roche Lightcycler using standard *reverse transcriptase* kit and SYBR Green Real-Time PCR Master Mixes (SIGMA).

### RNA-Seq: Mapping and quantification (Fig. 2)

Raw reads were mapped either with the Tophat pipeline or hisat2+stringtie pipeline due to logistic reasons, and is recorded in the supplementary TPM table. We used “Bdistachyon_314_v3.1” from Phytozome (*1*) as reference genome throughout the study.

**Hisat2+Stringtie** pipeline: Adapters were trimmed off from raw reads with Trimmomatic (*2*) with argument “ILLUMINACLIP:${{FA_ADAPTER}}:6:30:10 LEADING:3 TRAILING:3 MINLEN:36 SLIDINGWINDOW:4:15”. Raw reads were mapped to the transcriptome using HISAT2 (*3*) with argument:“--no-mixed --rna-strandness RF --dta --fr”. Duplicate reads were removed with Picard (*4*) using default setting. Transcripts were quantified according to this alignment with StringTie (*5*) in TPM values (Transcripts per Million mapped transcripts) with argument “--rf”.

**Tophat** pipeline

The TPM values were transformed into log abundances through

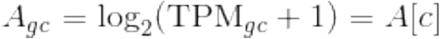

(*g* indexes genes while *c* indexes conditions).

Any gene with a maximum log abundance smaller than 3.0 was discarded from downstream analysis to avoid introducing noisy variation. Detailed TPM table can be found in the supplementary.

### RNA-Seq: Clustering and target calling

To investigate transcriptomic response towards a particular treatment, time-course perturbation matrices were constructed as the difference of log abundances between paired conditions. For example,

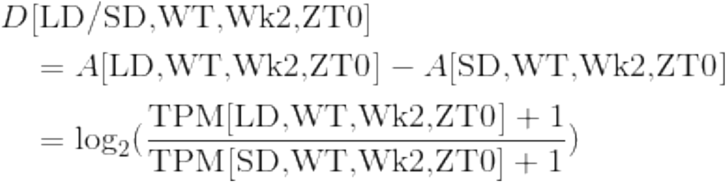

Clustering: The selected perturbation matrices were concatenated into a data matrix, against which a von Mises-Fisher mixture of increasing concentration is fitted (similar to clusterVMF (*6*)). Optimal concentration is manually selected to be ∼3.080 according to the diagnostic plot and clusters with an average uncertainty higher than 2.5 were considered non-significant.

The following perturbation matrices were selected:

- [LD/SD, WT, Wk2]
- [LD/SD, WT, Wk3]
- [SD, elf3-1/WT,Wk2]
- [SD, elf3-1/WT,Wk3]
- [LD, phyC-4/WT,Wk2]
- [LD, ppd1-1/WT,Wk3]

Selection of genes responsive to *elf3-1* and *phyC-4*: A continous timecourse function was interpolated linearly from each of the perturbation matrices [SD, *elf3-1*/WT,Wk2] and [LD, *phyC-4*/WT,Wk2], which is then integrated over 24 h to give a temporal average, 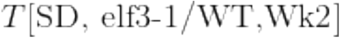 and 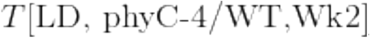 Genes that satisfy 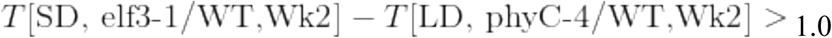 were considered transcriptionally perturbed.

### ChIP-seq experimental details (Fig. 3 and 4)

Seeds were sterilized and sown on ½ X Murashige and Skoog-agar (MS-agar) plates at pH 5.7 and grown in Magenta™ GA-7 Plant Culture Box (Thomas scientific) and harvested at the indicated time.

3 g seedlings for each set were fixed under vacuum for 20 min in 1xPBS (10 mM PO_4_^3−^, 137 mM NaCl, and 2.7 mM KCl) containing 1% Formaldehyde (F8775 SIGMA). The reaction was quenched by adding glycine to a final concentration of 62 mM. Chromatin immunoprecipitation (ChIP) was performed as described (*7*), with the exception that 100 µl FLAG M2 agarose affinity gel antibody was used (SIGMA-Aldrich) per sample. Sequencing libraries were prepared using TruSeq ChIP Sample Preparation Kit (Illumina IP-202-1024) and samples sequenced on NextSeq500 machine from Illumina using NextSeq® 500/550 High Output Kit v2 (75 cycles) TG-160-2005. Sequence reads were analysed using in-house pipelines.

### ChIP-Seq: Mapping and quantification

Adapters were trimmed off from raw reads with Trimmomatic (*2*) with argument “ILLUMINACLIP:${{FA_ADAPTER}}:6:30:10 LEADING:3 TRAILING:3 MINLEN:36 SLIDINGWINDOW:4:15”. Raw reads were mapped to the genome “Bdistachyon_314_v3.1” with Bowtie2 (*7*) under argument:“--no-mixed --no-discordant --no-unal -k2”. Any read that mapped to more than one genomic location was discarded. Duplicate reads were removed with Picard using default setting (*4*).

Genomic binding profile was quantified in RPKM (Reads Per Kilobase per Million mapped reads) using a bin-size of 10bp. For each bin,

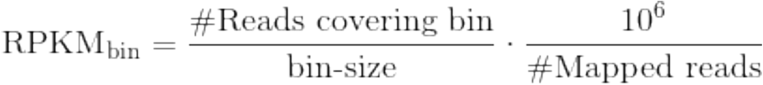

For each treated ChIP-Seq library, peaks were called against a control using MACS2 (*8*) with argument “--keep-dup 1 -p 0.1”. Peaks from all ChIP-Seqs were fitlered for FC>2.0.

### ChIP-Seq: Defining regions differentially bound by ELF3

Peaks from all ELF3 ChIP-Seqs were filtered and merged into contiguous regions. Abundances were computed for 100bp windows within each merged region, and log-transformed with

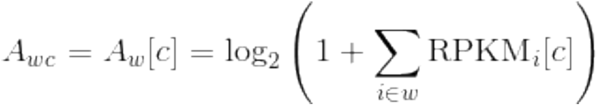

(*w* denotes the window, *i* denotes the 10bp bin, *c* indexes condition)

Differential occupancy was quantified for each window by taking its dot-product with a signature vector to get 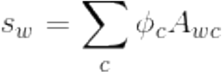, where the signature vector *ϕ_c_* is computed from a set of marker windows *W* so that

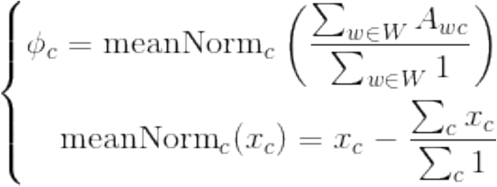

Cross-condition variance is also defined for each window 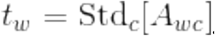. Any window with *s_g_* > 1.5 and *t_g_* > 0.5 is called differentially bound.

Any peak with a differentially bound window is considered a differentially-bound peak. Any gene with a differential bound window within +/− 6000bp of its start codon is considered a differentially bound gene. Pile-up of ELF3 occupancy: was performed on the set of peaks that contains at least 1 differentially bound window.

### ChIP-Seq: Peak-level overlap

Each ELF3 peak was considered overlapped if it’s within 1000bp of a phyC peak, and vice versa. The venn is constructed by specifying the number of non-overlapping ELF3 and phyC peaks, and the number of intersection peaks as

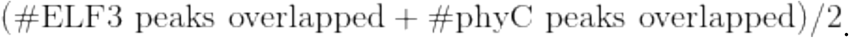

### Availability

Code is available from <u>Github repo</u>.

RNA-Seq data are available from: {{GEO_RNA}}

ChIP-Seq data are available from: {{GEO_CHIP}}

GNU-parallel (*11*) was used in paralleling the computational analysis.

Binding profiles from ChIP-Seq near marker genes were visualised with *fluff* (*9*), and visualised with Integrated Genomic Browser during investigation (*10*).

### Assaying dark reversion rate for PHYC (Fig. 4)

#### Lines used

*B. distachyon*

- WT
- phyC-ox in WT background; *PUB2-PHYC-ox* 19-7 (homozygous)

*A. thaliana* expressing *BdPHYC* in a *phyAB* mutant background (plant 5)

#### Method

*B. distachyon* seeds were incubated between 2 sheets of wet filter paper for 2-3 days in darkness at 4 °C. After removal of the lemma, the seeds were plated on ½ MS agar supplemented with 5 µM Norflurazon to inhibit greening during the red light irradiation. The seedlings were grown for 6 days at 22 °C in darkness. In order to induce the degradation of BdphyA and BdphyB the seedlings were irradiated with constant red light (660 nm, 10 µmol m^−2^ s^−1^) for 16 h. Subsequently, the seedlings were transferred to darkness at 22 °C to monitor dark reversion of BdphyC. At time points 0, 4, 8 and 12 h after dark transfer relative levels of active BdphyC (Pfr/Ptot) were measured using a dual wavelength ratio spectrophotometer (Ratiospect) as described previously (*8*). The shoot parts of 5-7 *B. distachyon* seedlings were used per measurement. To inhibit oxidation, the seedlings were incubated for 20 min in ice-cold 50 mM DTT solution prior to the measurement.

*A. thaliana* seeds were sterilised before plating them on 4 layers of Whatman® filter paper saturated with 4.5 ml ddH_2_O. For sterilisation, the seeds were washed first shortly with 70 % ethanol and then twice with 100 % ethanol. The seeds were stratified for at least 2 days at 4 °C in darkness. To induce germination, the seeds were incubated during 4 to 8 h in white light at 22 °C. Subsequently, the seedlings were grown in darkness at 22°C for 4 days. Prior to Ratiospect measurements, the seedlings were irradiated for 20 min with constant red light (660 nm, 10 µmol m^−2^ s^−1^) to convert BdphyC into the active Pfr form. Afterwards the seedlings were transferred into darkness darkness at 22 °C to monitor dark reversion of BdphyC. At time points 0, 4, 8 and 12 h after red light irradiation relative levels of active BdphyC (Pfr/Ptot) were measured using a dual wavelength ratio spectrophotometer (Ratiospect) as described previously (*8*). 120-140 mg of *A. thaliana* seedlings (freshweight) were used per measurement.

#### Proteomics

Plant materials for affinity purification coupled with mass spectrometry (AP-MS) were made from Brachypodium plants expressing either pUBI-ELF3-GFP-FLAG or pMDC32-YFP-His_6_-3xFLAG (negative control) transgene. After stratification in dark at 4 °C for 4 days, transgenic Brachypodium plants were grown in a growth chamber with a photoperiod of 14 hours of light (200 umol·m^−2^·s^−1^) and 10 hours of darkness, at 24 °C during daytime and 18 °C at night. Leaves from 45-day-old (old) or 21-day-old (young) plants were harvested at ZT0 in darkness and flash frozen in liquid N_2_. The protein extraction was performed in darkness with dim green safety light. About 30 mg (for old plants sample and YFP negative control) or 10 mg (for young plants sample) of total protein were used for purification via FLAG immune-precipitation (we used 1.4 µg anti-FLAG antibody per 1 mg total protein), using the method as previously described (*6*, *7*). After elution with 3xFLAG free peptides, eluates were precipitated by 25% TCA at −20°C, pelleted and washed with ice-cold acetone. Pellets were dried using a speed vacuum and sent for mass spectrometry analysis, with the same processing protocol and filtering criteria as described previously (*8*). MS data were extracted and searched against Brachypodium database to identify each protein (Phytozome 12, https://phytozome.jgi.doe.gov/pz/portal.html). All proteins identified in YFP control were subtracted from the identifications and a curated list containing BdELF3 specific interactors was presented, showing names of their Arabidopsis homolog proteins. log file of *ChIP-Seq* 188CR:

~~~
S1:
64-LD-ZT0-ELF3-OX-RERUN_S1.bowtie2
56859885 reads; of these:
  56859885 (100.00%) were paired; of these:
    10994569 (19.34%) aligned concordantly 0 times
    24159948 (42.49%) aligned concordantly exactly 1 time
    21705368 (38.17%) aligned concordantly >1 times
80.66% overall alignment rate
S2:
64-SD-ZT0-ELF3-OX-RERUN_S2.bowtie2
32907407 reads; of these:
  32907407 (100.00%) were paired; of these:
    12224010 (37.15%) aligned concordantly 0 times
    11387902 (34.61%) aligned concordantly exactly 1 time
    9295495 (28.25%) aligned concordantly >1 times
62.85% overall alignment rate
S3:
64-LL-ZT0-ELF3-OX-RERUN_S3.bowtie2
47414081 reads; of these:
  47414081 (100.00%) were paired; of these:
    12398525 (26.15%) aligned concordantly 0 times
    18170203 (38.32%) aligned concordantly exactly 1 time
    16845353 (35.53%) aligned concordantly >1 times
73.85% overall alignment rate
S4:
64-LD-ZT4-ELF3-OX-RERUN_S4.bowtie2
44936828 reads; of these:
  44936828 (100.00%) were paired; of these:
    12788499 (28.46%) aligned concordantly 0 times
    18976870 (42.23%) aligned concordantly exactly 1 time
    13171459 (29.31%) aligned concordantly >1 times
71.54% overall alignment rate
S5:
64-SD-ZT4-ELF3-OX-RERUN_S5.bowtie2
44752906 reads; of these:
  44752906 (100.00%) were paired; of these:
    16478316 (36.82%) aligned concordantly 0 times
    15698163 (35.08%) aligned concordantly exactly 1 time
    12576427 (28.10%) aligned concordantly >1 times
63.18% overall alignment rate
S6:
66-LD-ZT4-PRR37OX-RERUN_S6.bowtie2
44752906 reads; of these:
  44752906 (100.00%) were paired; of these:
    16478316 (36.82%) aligned concordantly 0 times
    15698163 (35.08%) aligned concordantly exactly 1 time
    12576427 (28.10%) aligned concordantly >1 times
63.18% overall alignment rate
S7
64-LD-ZT20-ELF3OX-RERUN_S7.bowtie2
54058806 reads; of these:
  54058806 (100.00%) were paired; of these:
    15052008 (27.84%) aligned concordantly 0 times
    21878336 (40.47%) aligned concordantly exactly 1 time
    17128462 (31.68%) aligned concordantly >1 times
72.16% overall alignment rate
S8:
64-SD-ZT20-ELF3OX-RERUN_S8.bowtie2
63445898 reads; of these:
  63445898 (100.00%) were paired; of these:
    14483986 (22.83%) aligned concordantly 0 times
    28850417 (45.47%) aligned concordantly exactly 1 time
    20111495 (31.70%) aligned concordantly >1 times
77.17% overall alignment rate

S9:
phyC-OX-GFP-FLAG-Zt20-LD-RERUN_S9.bowtie2
49021059 reads; of these:
  49021059 (100.00%) were paired; of these:
    11516381 (23.49%) aligned concordantly 0 times
    17325692 (35.34%) aligned concordantly exactly 1 time
    20178986 (41.16%) aligned concordantly >1 times
76.51% overall alignment rate

S10:
phyC-OX-GFP-FLAG-Zt20-LD-RERUN_S10.bowtie2
43207042 reads; of these:
  43207042 (100.00%) were paired; of these:
    9431466 (21.83%) aligned concordantly 0 times
    17663422 (40.88%) aligned concordantly exactly 1 time
    16112154 (37.29%) aligned concordantly >1 times
78.17% overall alignment rate

175C run_INPUT-BD_S4
22572371 reads; of these:
  22572371 (100.00%) were unpaired; of these:
    1604341 (7.11%) aligned 0 times
    12073412 (53.49%) aligned exactly 1 time
    8894618 (39.40%) aligned >1 times
92.89% overall alignment rate
~~~

## References

Alvarez, M. A., Tranquilli, G., Lewis, S., Kippes, N., & Dubcovsky, J. (2016). Genetic and physical mapping of the earliness per se locus Eps-A m 1 in Triticum monococcum identifies EARLY FLOWERING 3 (ELF3) as a candidate gene. Functional & Integrative Genomics, 16(4), 365–382. https://doi.org/10.1007/s10142-016-0490-3

Borthwick, H. A., & Hendricks, S. B. (1960). Photoperiodism in Plants. Science (New York, N.Y.), 132(3435), 1223–1228. https://doi.org/10.1126/science.132.3435.1223

Campoli, C., Pankin, A., Drosse, B., Casao, C. M., Davis, S. J., & von Korff, M. (2013). *HvLUX1* is a candidate gene underlying the *early maturity 10* locus in barley: phylogeny, diversity, and interactions with the circadian clock and photoperiodic pathways. New Phytologist, 199(4), 1045–1059. https://doi.org/10.1111/nph.12346

Chen, A., Li, C., Hu, W., Lau, M. Y., Lin, H., Rockwell, N. C., … Dubcovsky, J. (2014). PHYTOCHROME C plays a major role in the acceleration of wheat flowering under long-day photoperiod. Proceedings of the National Academy of Sciences, 111(28), 10037–10044. https://doi.org/10.1073/pnas.1409795111

Cumming, B. G., Hendricks, S. B., & Borthwick, H. A. (1965). RHYTHMIC FLOWERING RESPONSES AND PHYTOCHROME CHANGES IN A SELECTION OF CHENOPODIUM RUBRUM. Canadian Journal of Botany, 43(7), 825–853. https://doi.org/10.1139/b65-092

Digel, B., Tavakol, E., Verderio, G., Tondelli, A., Xu, X., Cattivelli, L., … von Korff, M. (2016). Photoperiod-H1 (Ppd-H1) Controls Leaf Size. Plant Physiology, 172(1), 405–415. https://doi.org/10.1104/pp.16.00977

Ezer, D., Jung, J.-H., Lan, H., Biswas, S., Gregoire, L., Box, M. S., … Wigge, P. A. (2017). The evening complex coordinates environmental and endogenous signals in Arabidopsis. Nature Plants, 3(7), 17087. https://doi.org/10.1038/nplants.2017.87

Faure, S., Turner, A. S., Gruszka, D., Christodoulou, V., Davis, S. J., von Korff, M., & Laurie, D. A. (2012). Mutation at the circadian clock gene EARLY MATURITY 8 adapts domesticated barley (Hordeum vulgare) to short growing seasons. Proceedings of the National Academy of Sciences. https://doi.org/10.1073/pnas.1120496109

Goralogia, G. S., Liu, T.-K., Zhao, L., Panipinto, P. M., Groover, E. D., Bains, Y. S., & Imaizumi, T. (2017). CYCLING DOF FACTOR 1 represses transcription through the TOPLESS co-repressor to control photoperiodic flowering in Arabidopsis. The Plant Journal, 92(2), 244–262. https://doi.org/10.1111/tpj.13649

Hamner, K. C. (1940). Interrelation of Light and Darkness in Photoperiodic Induction. Botanical Gazette, 101(3), 658–687. https://doi.org/10.1086/334903

Hayama, R., Sarid-Krebs, L., Richter, R., Fernández, V., Jang, S., & Coupland, G. (2017). PSEUDO RESPONSE REGULATORs stabilize CONSTANS protein to promote flowering in response to day length. The EMBO Journal. https://doi.org/10.15252/embj.201693907

Huang, H., Gehan, M. A., Huss, S. E., Alvarez, S., Lizarraga, C., Gruebbling, E. L., … Nusinow, D. A. (2017). Cross-species complementation reveals conserved functions for EARLY FLOWERING 3 between monocots and dicots. Plant Direct, 1(4), e00018. https://doi.org/10.1002/pld3.18

Hughes, J. E., Morgan, D. C., Lambton, P. A., Black, C. R., & Smith, H. (1984). Photoperiodic time signals during twilight. Plant, Cell and Environment, 7(4), 269–277. https://doi.org/10.1111/1365-3040.ep11589464

Ishikawa, R., Shinomura, T., Takano, M., & Shimamoto, K. (2009). Phytochrome dependent quantitative control of Hd3a transcription is the basis of the night break effect in rice flowering. Genes & Genetic Systems, 84(2), 179–184. Retrieved from http://www.ncbi.nlm.nih.gov/pubmed/19556711

Ishikawa, R., Tamaki, S., Yokoi, S., Inagaki, N., Shinomura, T., Takano, M., & Shimamoto, K. (2005). Suppression of the floral activator Hd3a is the principal cause of the night break effect in rice. Plant Cell, 17(12), 3326–3336. Retrieved from http://www.ncbi.nlm.nih.gov/entrez/query.fcgi?cmd=Retrieve&db=PubMed&dopt=Citation&list_uids=16272430

Itoh, H., Tanaka, Y., & Izawa, T. (2018). Genetic relationship between phytochromes and *OsELF3-1* reveals the mode of regulation for the suppression of phytochrome signaling in rice. Plant and Cell Physiology. https://doi.org/10.1093/pcp/pcy225

Jos Ramos-Sánchez, A. M., Triozzi, P. M., Alique, D., Wigge, P. A., Allona, I., & Perales, M. (2019). LHY2 Integrates Night-Length Information to Determine Timing of Poplar Photoperiodic Growth. Current Biology, 29. https://doi.org/10.1016/j.cub.2019.06.003

Jung, J.-H., Domijan, M., Klose, C., Biswas, S., Ezer, D., Gao, M., … Wigge, P. A. (2016). Phytochromes function as thermosensors in Arabidopsis. Science, 354(6314).

Legris, M., Klose, C., Burgie, E. S., Rojas, C. C. R., Neme, M., Hiltbrunner, A., … Casal, J. J. (2016). Phytochrome B integrates light and temperature signals in Arabidopsis. Science, 354(6314).

Linkosalo, T., & Lechowicz, M. J. (2006). Twilight far-red treatment advances leaf bud burst of silver birch (Betula pendula). Tree Physiology, 26(10), 1249–1256. Retrieved from http://www.ncbi.nlm.nih.gov/pubmed/16815827

Mizuno, N., Kinoshita, M., Kinoshita, S., Nishida, H., Fujita, M., Kato, K., … Nasuda, S. (2016). Loss-of-Function Mutations in Three Homoeologous PHYTOCLOCK 1 Genes in Common Wheat Are Associated with the Extra-Early Flowering Phenotype. PLOS ONE, 11(10), e0165618. https://doi.org/10.1371/journal.pone.0165618

Nanda, K. K., & Hamner, K. C. (1958). Studies on the Nature of the Endogenous Rhythm Affecting Photoperiodic Response of Biloxi Soybean. Botanical Gazette, 120(1), 14–25. https://doi.org/10.1086/335992

Nishida, H., Ishihara, D., Ishii, M., Kaneko, T., Kawahigashi, H., Akashi, Y., … Kato, K. (2013). Phytochrome C Is A Key Factor Controlling Long-Day Flowering in Barley. PLANT PHYSIOLOGY, 163(2), 804–814. https://doi.org/10.1104/pp.113.222570

Olsen, J. E., Junttila, O., Nilsen, J., Eriksson, M. E., Martinussen, I., Olsson, O., … Moritz, T. (1997). Ectopic expression of oat phytochrome A in hybrid aspen changes critical daylength for growth and prevents cold acclimatization. The Plant Journal, 12(6), 1339–1350. https://doi.org/10.1046/j.1365-313x.1997.12061339.x

Pankin, A., Campoli, C., Dong, X., Kilian, B., Sharma, R., Himmelbach, A., … von Korff, M. (2014). Mapping-by-sequencing identifies HvPHYTOCHROME C as a candidate gene for the early maturity 5 locus modulating the circadian clock and photoperiodic flowering in barley. Genetics, 198(1), 383–396. https://doi.org/10.1534/genetics.114.165613

Pearce, S., Shaw, L. M., Lin, H., Cotter, J. D., Li, C., & Dubcovsky, J. (2017). Night-Break Experiments Shed Light on the Photoperiod1-Mediated Flowering. Plant Physiology, 174(2), 1139–1150. https://doi.org/10.1104/pp.17.00361

Roden, L. C., Song, H.-R., Jackson, S., Morris, K., & Carre, I. A. (2002). Floral responses to photoperiod are correlated with the timing of rhythmic expression relative to dawn and dusk in Arabidopsis. Proceedings of the National Academy of Sciences of the United States of America, 99(20), 13313–13318. https://doi.org/10.1073/pnas.192365599

Rubenach, A. J. S., Hecht, V., Vander Schoor, J. K., Liew, L. C., Aubert, G., Burstin, J., & Weller, J. L. (2017). EARLY FLOWERING3 Redundancy Fine-Tunes Photoperiod Sensitivity. Plant Physiology, 173(4), 2253–2264. https://doi.org/10.1104/pp.16.01738

Seki, M., Chono, M., Nishimura, T., Sato, M., Yoshimura, Y., Matsunaka, H., … Kato, K. (2013). Distribution of photoperiod-insensitive allele Ppd-A1a and its effect on heading time in Japanese wheat cultivars. Breeding Science, 63(3), 309–316. https://doi.org/10.1270/jsbbs.63.309

Song, Y. H., Shim, J. S., Kinmonth-Schultz, H. A., & Imaizumi, T. (2015). Photoperiodic flowering: time measurement mechanisms in leaves. Annual Review of Plant Biology, 66, 441–464. https://doi.org/10.1146/annurev-arplant-043014-115555

Thomas, B., & Vince-Prue, D. (1997). Photoperiodism in Plants. London: Academic Press.

Turner, A., Beales, J., Faure, S., Dunford, R. P., & Laurie, D. A. (2005). The pseudo-response regulator Ppd-H1 provides adaptation to photoperiod in barley. Science, 310(5750), 1031–1034. https://doi.org/10.1126/science.1117619

Wigge, P. A. (2011). FT, a mobile developmental signal in plants. Current Biology: CB, 21(9), R374–8. https://doi.org/10.1016/j.cub.2011.03.038

Wilhelm, E. P., Turner, A. S., & Laurie, D. A. (2009). Photoperiod insensitive Ppd-A1a mutations in tetraploid wheat (Triticum durum Desf.). Theoretical and Applied Genetics, 118(2), 285–294. https://doi.org/10.1007/s00122-008-0898-9

Woods, D. P., Ream, T. S., Minevich, G., Hobert, O., & Amasino, R. M. (2014). PHYTOCHROME C Is an Essential Light Receptor for Photoperiodic Flowering in the Temperate Grass, Brachypodium distachyon. Genetics, genetics.114.166785-. https://doi.org/10.1534/genetics.114.166785

## REFERENCES

1. D. P. Woods, T. S. Ream, G. Minevich, O. Hobert, R. M. Amasino, Genetics, in press, doi:10.1534/genetics.114.166785.

2. J. Miao et al., Targeted mutagenesis in rice using CRISPR-Cas system. Cell Res. 23, 1233–6 (2013).

3. S. C. Alves et al., A protocol for Agrobacterium-mediated transformation of Brachypodium distachyon community standard line Bd21. Nat Protoc. 4, 638–649 (2009).

4. H. Huang et al., PCH1 integrates circadian and light-signaling pathways to control photoperiod-responsive growth in Arabidopsis. Elife. 5 (2016), doi:10.7554/eLife.13292.

5. M. D. Curtis, U. Grossniklaus, A Gateway Cloning Vector Set for High-Throughput Functional Analysis of Genes in Planta. PLANT Physiol. 133, 462–469 (2003).

6. J. Vogel, T. Hill, High-efficiency Agrobacterium-mediated transformation of Brachypodium distachyon inbred line Bd21-3. Plant Cell Rep. 27, 471–478 (2008).

7. K. E. Jaeger, N. Pullen, S. Lamzin, R. J. Morris, P. A. Wigge, Interlocking feedback loops govern the dynamic behavior of the floral transition in Arabidopsis. Plant Cell. 25, 820–33 (2013).

8. J. Rausenberger et al., An integrative model for phytochrome B mediated photomorphogenesis: From protein dynamics to physiology. PLoS One. 5 (2010), doi:10.1371/journal.pone.0010721.

9. H. Huang, D. Nusinow, Tandem Purification of His6-3x FLAG Tagged Proteins for Mass Spectrometry from Arabidopsis. BIO-PROTOCOL. 6 (2016), doi:10.21769/BioProtoc.2060.

10. H. Huang et al., Identification of Evening Complex Associated Proteins in Arabidopsis by Affinity Purification and Mass Spectrometry. Mol. Cell. Proteomics. 15, 201–217 (2016).

11. J. A. Vizcaíno et al., ProteomeXchange provides globally coordinated proteomics data submission and dissemination. Nat. Biotechnol. 32, 223–6 (2014).

